# Accurate variant effect estimation in FACS-based deep mutational scanning data with Lilace

**DOI:** 10.1101/2025.06.24.661380

**Authors:** Jerome Freudenberg, Jingyou Rao, Matthew K. Howard, Christian Macdonald, Noah F. Greenwald, Willow Coyote-Maestas, Harold Pimentel

## Abstract

Deep mutational scanning (DMS) experiments interrogate the effect of genetic variants on protein function, often using fluorescence-activated cell sorting (FACS) to quantitatively measure molecular phenotypes, such as abundance or activity. Analysis of DMS experiments with a FACS readout is challenging due to measurement variance and the unique multidimensional nature of the phenotype. However, no statistical method has yet been developed to address the challenges of FACS-based DMS. Here we present Lilace, a Bayesian statistical model to estimate variant effects with uncertainty quantification from FACS-based DMS experiments. We validate Lilace’s performance and robustness using simulated data and apply it to OCT1 and Kir2.1 DMS experiments, demonstrating an improved false discovery rate (FDR) while largely maintaining sensitivity.

## Introduction

Deep mutational scanning (DMS) experiments help elucidate sequence-function relationships by systematically interrogating the full spectrum of amino acid substitution effects on the molecular properties and function of a protein (1; 2; 3; 4). They provide insight into the mechanisms of protein function and variation (5; 6), as well as aid clinical interpretation of genetic variants (7; 8; 9). When coupled with fluorescence-activated cell sorting (FACS), DMS can scalably measure a quantitative phenotype across variants (1; 10), such as protein abundance (10; 11; 12; 13; 14; 15), cell surface expression (12; 16; 17; 18), and activity (11; 17; 18). In such FACS-based DMS experiments, a pooled library of variants is sorted into bins based on fluorescence intensity; then, each bin is sequenced (Fig. 1A). The resulting read counts act as a proxy for the proportion of variants in each bin, representing the tagged phenotype distribution of each variant. To distinguish between variants with approximately wild type-like behavior and those with clear impacts on the phenotype, each variant is compared to that of a negative control group, such as synonymous variants (13; 14; 19). To facilitate this comparison, variants are scored on a quantitative scale based on the difference between their phenotype distribution and the negative control phenotype.

**Figure 1:**
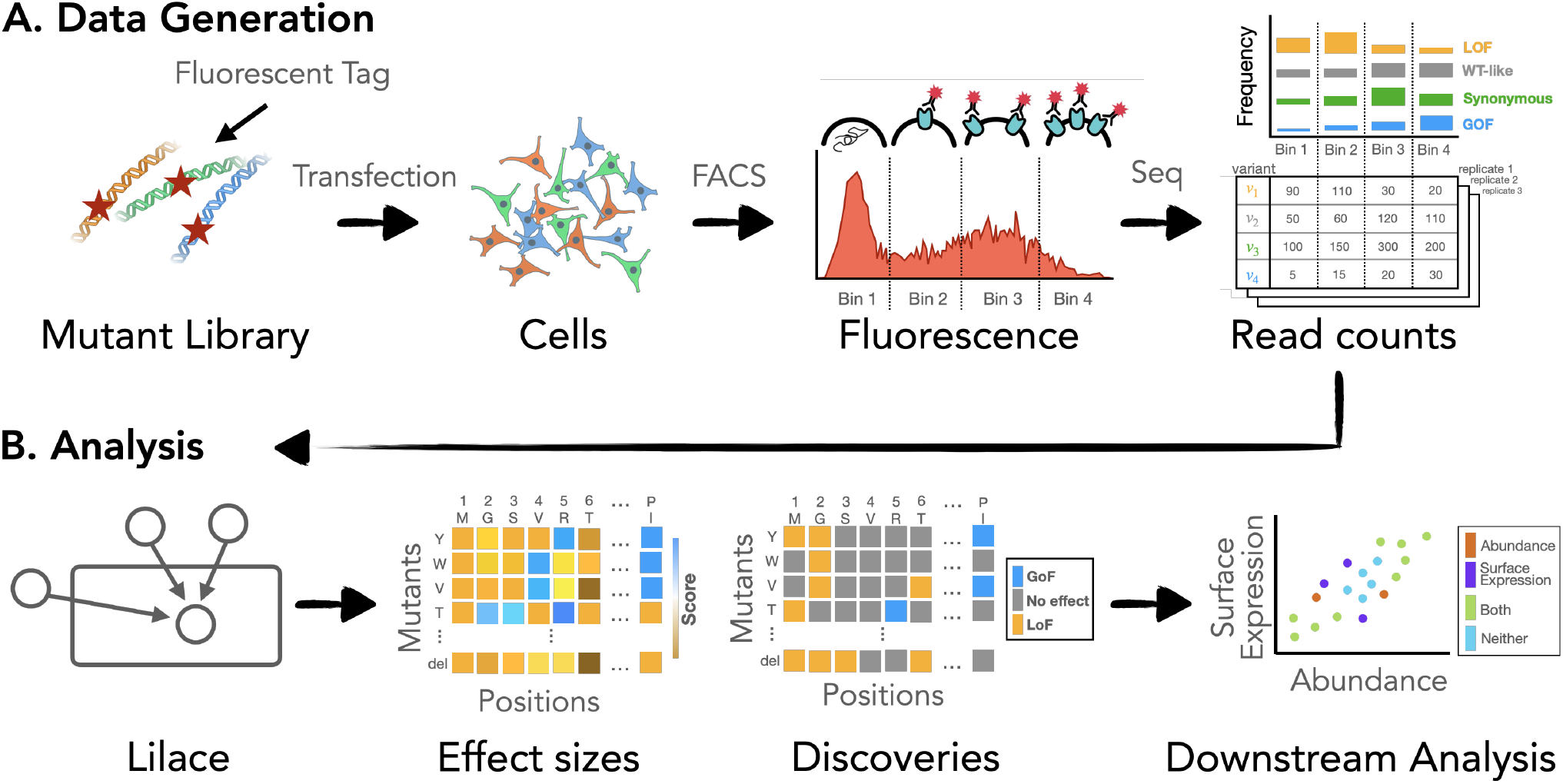
(A) Overview of workflow from experimental process to Lilace input. (B) Uncertainty quantification enables analysis based on statistical significance in addition to effect size.

A highly reproducible and accurate set of scores is essential for ranking high-effect variants to pinpoint functionally relevant protein regions. Additionally, estimating score uncertainty from replicates enables reliability assessment of individual variant scores for potential downstream and clinical interpretation (8; 20). However, scoring variants based on their FACS readout presents unique statistical challenges, including low sample sizes (two to three replicates is common), experimental noise, and low resolution of the fluorescence distribution due to limited FACS bins (with a standard of four bins). A common scoring approach is to compute a count-weighted mean of integer-labeled bins for each variant (10; 11; 15), then apply a significance cut-off based on the standard deviation of the synonymous variant score distribution (13; 14; 17). However, determining significance based on a variant’s mean score ignores variability between replicates and does not perform an explicit hypothesis test (i.e. p-value or analogous test-statistic). The lack of an explicit test results in ad hoc choice of the significance threshold and precludes control of the false discovery rate (FDR), which is necessary to produce reliable results. Another previously used scoring approach applies statistical models designed for growth-based DMS experiments to incorporate measurement uncertainty (14; 17; 20; 21). However, growth-based modeling assumptions, such as a consistent slope across a variant’s bin counts, are misspecified for the FACS readout. In practice, the uncertainty estimates from these models are not used for FACS data (14; 17; 21).

Here we present Lilace, a Bayesian hierarchical model developed specifically for FACS-based DMS data (Fig. 1) that inherently solves the problem of choosing a discovery threshold while providing reproducible and accurate results. The model estimates variant effects relative to a negative control baseline and quantifies effect uncertainties using replicate variance and a negative control-based bias correction. Lilace builds on the efficacy of Rosace (22), an analogous model for growth-based DMS data, and waterbear (23), a model for FACS-based pooled CRISPR screens. Similarly to Rosace, we incorporate a positional effect and a count-variance relationship to improve power in small sample sizes. Similarly to waterbear, we model a latent unimodal shift in the fluorescence distribution to filter experimental noise. We additionally introduce a data normalization procedure to account for disproportionate PCR amplification bias when the FACS gates are not set on equal proportions of the overall fluorescence distribution.

To validate Lilace’s performance, we simulated a wide range of experimental designs to quantify and compare discovery set accuracy with previous approaches, which, to our knowledge, we are also the first to benchmark. We simulated from a more complex generative model of the data, inspired by previous FACS simulation designs (23; 24), using simulation parameters estimated from existing datasets. We found that Lilace provided a lower false discovery rate (FDR) across a variety of simulations, while mostly maintaining sensitivity outside of high experimental variance scenarios. In these high-variance simulations, Lilace was the only method to retain a low, consistent FDR and, nevertheless, still retained some sensitivity. We further investigated the performance and generalizability of Lilace by computing empirical performance metrics on real datasets from different experimental procedures, with broad agreement with our simulations. We also examined how Lilace’s uncertainty calculation changed the discovery sets in previous screens of human organic cation transporter OCT1 (14) and inward rectifier potassium channel Kir2.1 (17), corresponding to better agreement with prior biological expectations.

## Results

### Hierarchical Bayesian model for FACS-based DMS data

To address the difficulties with statistical inference in FACS-based DMS data, we constructed Lilace to capture the structure of the data in the structure of the model (Fig. 2). Lilace builds on three core ideas: a variant’s effect is manifested as a shift in a unimodal fluorescence distribution, a variant’s count is a noisy observation of the underlying fluorescence distribution, and a variant tends to have similar effects as others at the same residue position.

**Figure 2:**
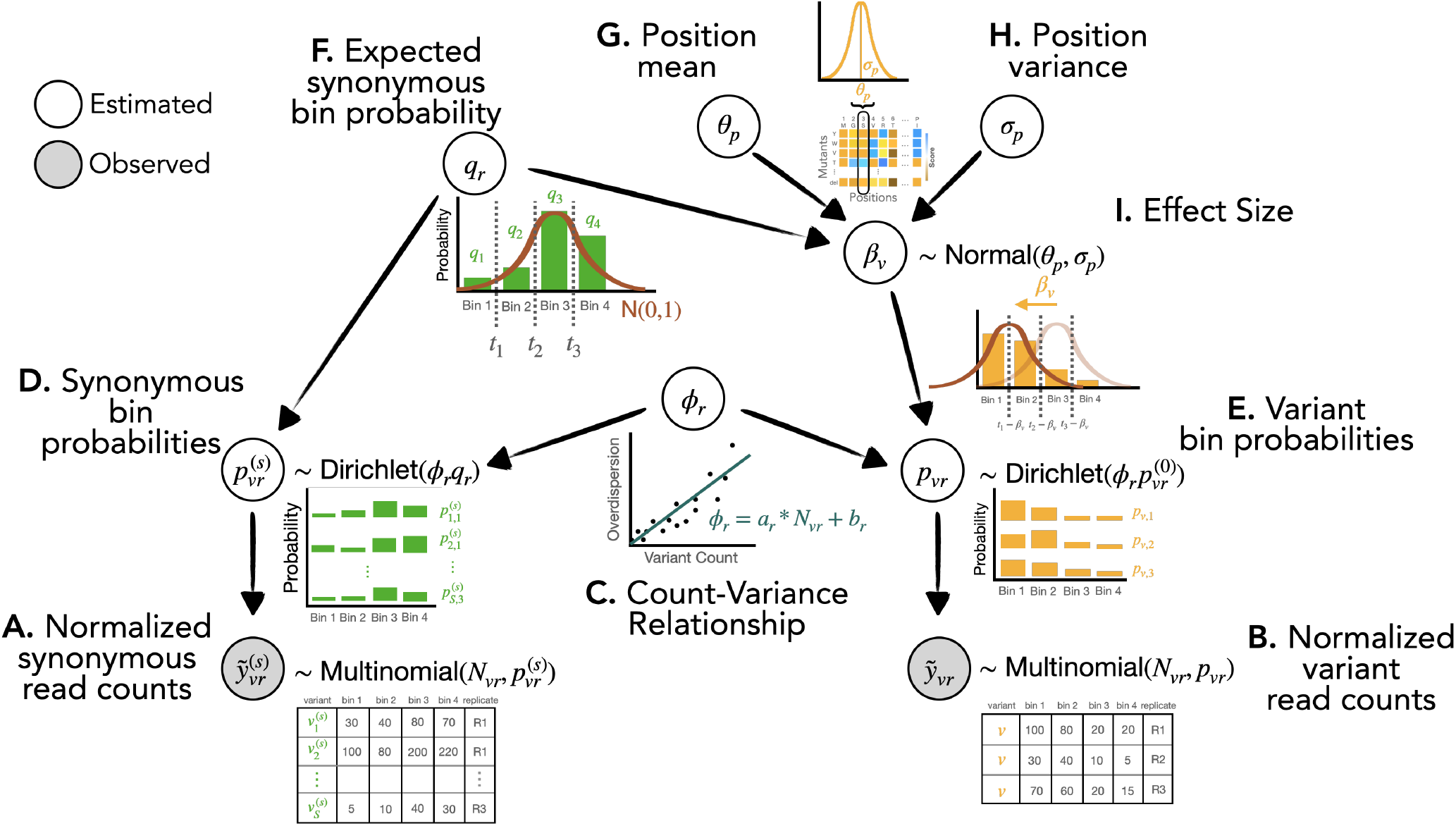
Conceptual visualization of generative model hierarchy. (A,B) Observed read counts are normalized to cell sorting populations, if available. (C) Overdispersion in read counts is modeled with a count-overdispersion trend function. (D,E) Each variant’s read counts are modeled with a latent probability of observing a count in each bin. (F) The expected probability of the negative control group (synonymous variants) within a replicate serves as a baseline to score against. (G,H) Variant effect sizes are informed by their position’s score distribution. (I) The effect size is modeled as a shift in the latent bin probabilities between a variant and the negative control baseline.

Fundamentally, Lilace places a constraint on the nature of possible changes between individual variant fluorescence distributions, modeling effect sizes as a unimodal shift from the negative control distribution (Fig. 2I, Supp Fig. 2A). While we use synonymous variants as our negative controls, this group could also, in principle, be composed of a different category of variants, such as a true wildtype. More specifically, Lilace models the bin count proportions for a variant v using a latent replicate-specific bin probability 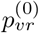 (Fig. 2E). The effect size for variant *v* is then modeled as the difference between 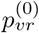 and the replicate-specific expected bin probability of synonymous variants *q*_*r*_ (Fig. 2F,I). The bin probabilities 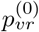 and *q*_*r*_ are mapped onto corresponding quantile cut-offs in a standard normal and the effect size is fitted as the distance between the cut-offs (Methods). This latent standard normal mapping constrains the possible estimated bin proportions of a variant’s fluorescence distribution to match a unimodal shift from the synonymous fluorescence distribution. Count proportion plots of DMS datasets show that such an effect size trajectory is a reasonable assumption (Supp Fig. 2B).

Since the observed counts are a noisy representation of the underlying fluorescence distribution, Lilace additionally incorporates an overdispersion parameter *ϕ* to capture this variance, modeled with a count-variance trend function (Fig. 2C). Such count-variance functions are commonly modeled in biological count data (22; 25)–in the case of FACS-based DMS data, we find a linear model to be sufficient (Supp Fig. 17). Lilace also incorporates the residual variance in synonymous variant scores into the effect size uncertainty as a negative control-based bias correction (26) (Methods). We expect synonymous variants to display wild type-like behavior (27), so we can use the variance in their scores to help distinguish variants between wild type-like and those with sufficient evidence of an effect.

Lastly, Lilace incorporates positional information to regularize effect estimates using a position-specific mean and variance (Fig. 2G,H). Sharing information within a position via this Bayesian hierarchy was shown by Rao et al. (2024) to improve sensitivity in DMS datasets (22). While we expect modeling position-level effects to improve estimation for most experiments, as observed in Rosace (22), we also provide the option to run Lilace without a position hierarchy for positionally unbiased estimates. Additionally, to avoid excessive shrinkage in the synonymous mutation effect sizes and preserve their ability to act as negative controls, Lilace incorporates a fixed effect for each synonymous mutation with a single shared variance term. This approach keeps the relative ranking of their scores consistent with their count observations among all variants (Supp Fig. 16).

### Lilace has improved FDR with comparable sensitivity

To examine our model’s performance against a known ground truth, we developed simulations informed by marginal parameter distributions from real data (Methods). We set default simulation parameters based on parameter estimates from an input dataset, then toggled each parameter individually to examine the effect of different sources of dataset heterogeneity on model performance (Fig. 3A). As input datasets for our simulation, we used an abundance screen on OCT1, performed with a protocol based on VAMP-seq (10), and a surface expression screen on Kir2.1, performed with a fluorescent antibody tagging protocol (12). Using our simulated ground truth, we then benchmarked the false discovery rate (FDR) and sensitivity of Lilace with previously applied approaches, including the most widely used growth-based DMS tool Enrich2 (20), a custom maximum likelihood-based scoring approach based on previous Sort-seq estimators (28), and a weighted bin average (10; 11) that uses the synonymous variant score distribution to determine a discovery threshold. The latter two act as a general proxy for methods that attempt to either estimate the actual variant fluorescence or non-parametrically score variants using a mean of integer-labeled bins, since implementations can vary in practice.

**Figure 3:**
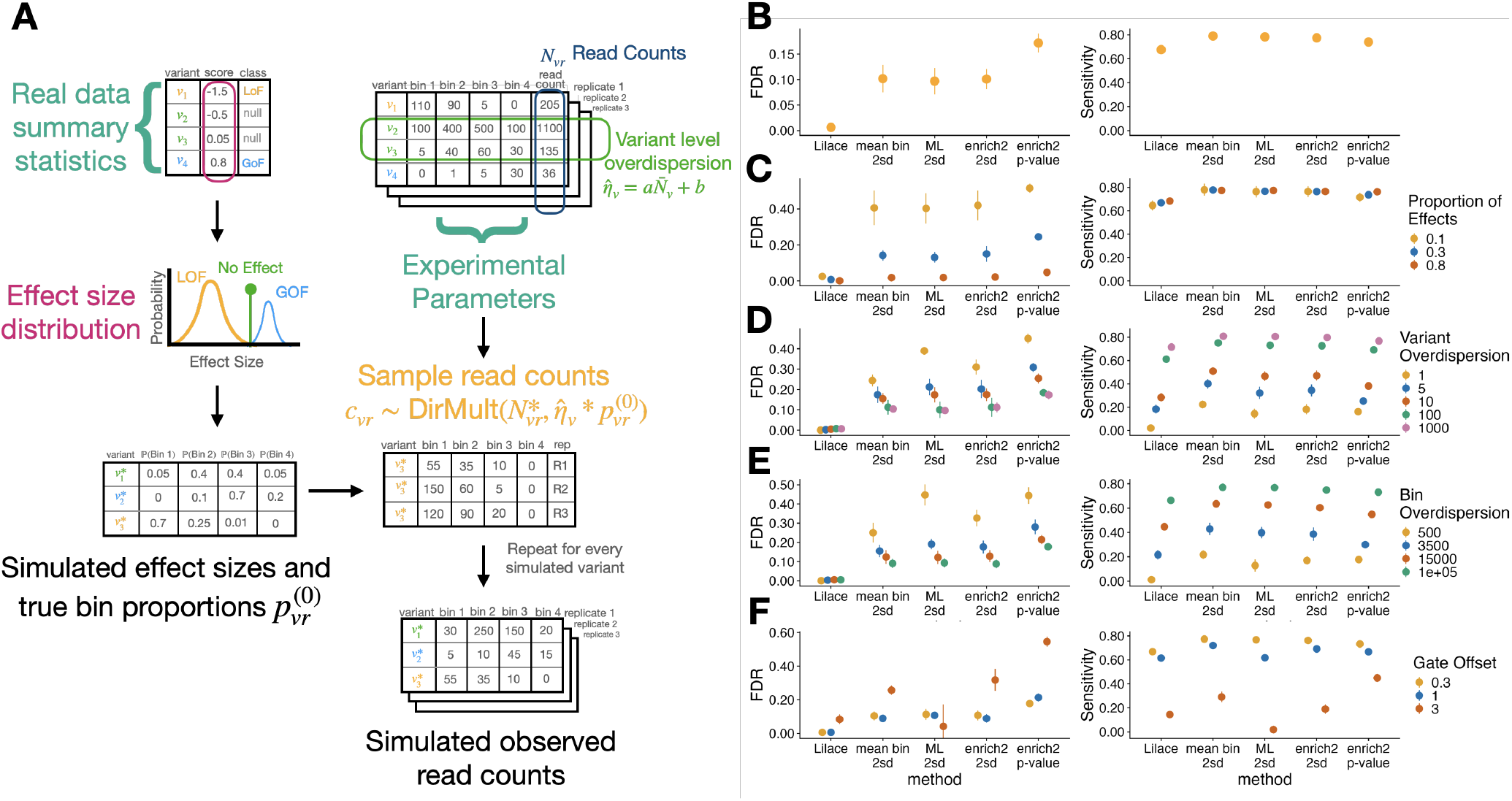
(A) Conceptual visualization of generative model used for simulations. The effect distribution and experimental noise parameters for the simulation were estimated from an input dataset. (B-F) FDR and sensitivity metrics from OCT1 abundance-seeded simulations. Plots include the (B) data-estimated default setting and toggling various parameters, including (C) proportion of variants with an effect, (D) variant overdispersion, (E) bin overdispersion, and (F) gate offset. Standard error bars are computed over ten iterations initialized with different random seeds. Remaining parameter figures and Kir2.1 surface expression-seeded simulation results are included in supplementary figures 3 and 4.

In the data-estimated default setting, we found that Lilace achieved the lowest FDR, with around a 95% reduction from the next best performing method (Fig. 3B, Supp Figs. 3, 4). Lilace continued to provide better FDR across simulation configurations resulting from both biological and experimental variability (Fig. 3C-F, Supp Figs. 3, 4). This improvement in FDR was especially pronounced in configurations with a low proportion of variants with an effect (Fig. 3C), high variance between individual variant observations (Fig. 3D), or bin sequencing variance (Fig. 3E).

Lilace generally retained power to detect effects, but prioritized FDR control, resulting in marginally lower sensitivity. In the default simulation, Lilace had slightly lower sensitivity than the highest sensitivity approaches (Fig. 3B, Supp Fig. 3, 4). We suspected this was driven by the large proportion of small effects in the default simulation (Methods). Filtering to the top 50% of effects mitigated this decrease, supporting this hypothesis and indicating that Lilace remained well-powered to detect more impactful variants (Supp Fig. 5). In the harder simulation configurations, such as a small variant-specific overdispersion parameter, Lilace was the only approach with a low FDR, at the cost of substantially lower sensitivity than other approaches (Fig. 3D). We further explored this FDR-sensitivity trade-off across approaches by examining discovery threshold-agnostic performance. For example, setting the mean bin discovery threshold to greater synonymous standard deviations would also lead to a reduction in sensitivity in favor of lower FDR. We found that while Lilace performed the best in this metric, most approaches had a similar overall trade-off (Supp Fig. 6). However, a main strength of Lilace lies in adaptively adjusting discovery classifications at a 95% certainty cut-off to keep FDR consistent in noisy data, whereas fixed discovery threshold approaches do not have this flexibility. As a result, Lilace is more robust to sources of experimental variance.

We also benchmarked performance across FACS-related gating parameters, including the number of bins and a gate offset parameter (Fig. 3F). In our simulation defaults, we set FACS gates to contain an equal proportion of the overall fluorescence distribution; however, gating can be done differently, especially when examining multiple conditions (21). For example, gating thresholds could be determined in an inactive protein state, shifting the fluorescence distribution of the active protein toward the rightmost bins. We found that severely shifting the gates from the equal proportion default hurt performance for all effect estimation approaches (Fig. 3F). This behavior is expected, since differentiating effects is harder if they are manifested within a smaller range of bins. Most approaches had a resulting increase in FDR and a decrease in sensitivity, while Lilace again retained a consistently low FDR.

In addition to simulations, we examined a more direct empirical proxy for FDR in real datasets by randomly masking 20% of synonymous mutations, treating them as any other variants, and checking how many are called incorrectly. We applied this analysis to a range of previously published FACS-based screens to benchmark generalizability, including OCT1, Kir2.1, GPR68, PTEN, TPMT, and P2RY8 (10; 12; 14; 18; 21). We found that Lilace again provided the best FDR using this metric (Fig. 4A). Since there is no analogous positive control set for high-effect variants, we instead used AlphaMissense pathogenicity predictions as an imperfect proxy for tracking relative method performance (29). We found that Lilace had a lower FDR derived from benign predictions and comparable sensitivity derived from pathogenic predictions, affirming our simulation conclusions (Fig. 4B). Lilace effect sizes also had the highest or equivalent to the highest correlation with the AlphaMissense pathogenicity scores for every dataset tested (Supp Table 1), suggesting that Lilace can reduce noise in variant scores. We also empirically assessed sensitivity using nonsense variants and ClinVar pathogenic-labeled variants, where available (Supp Fig. 8) (30), with similar results.

**Figure 4:**
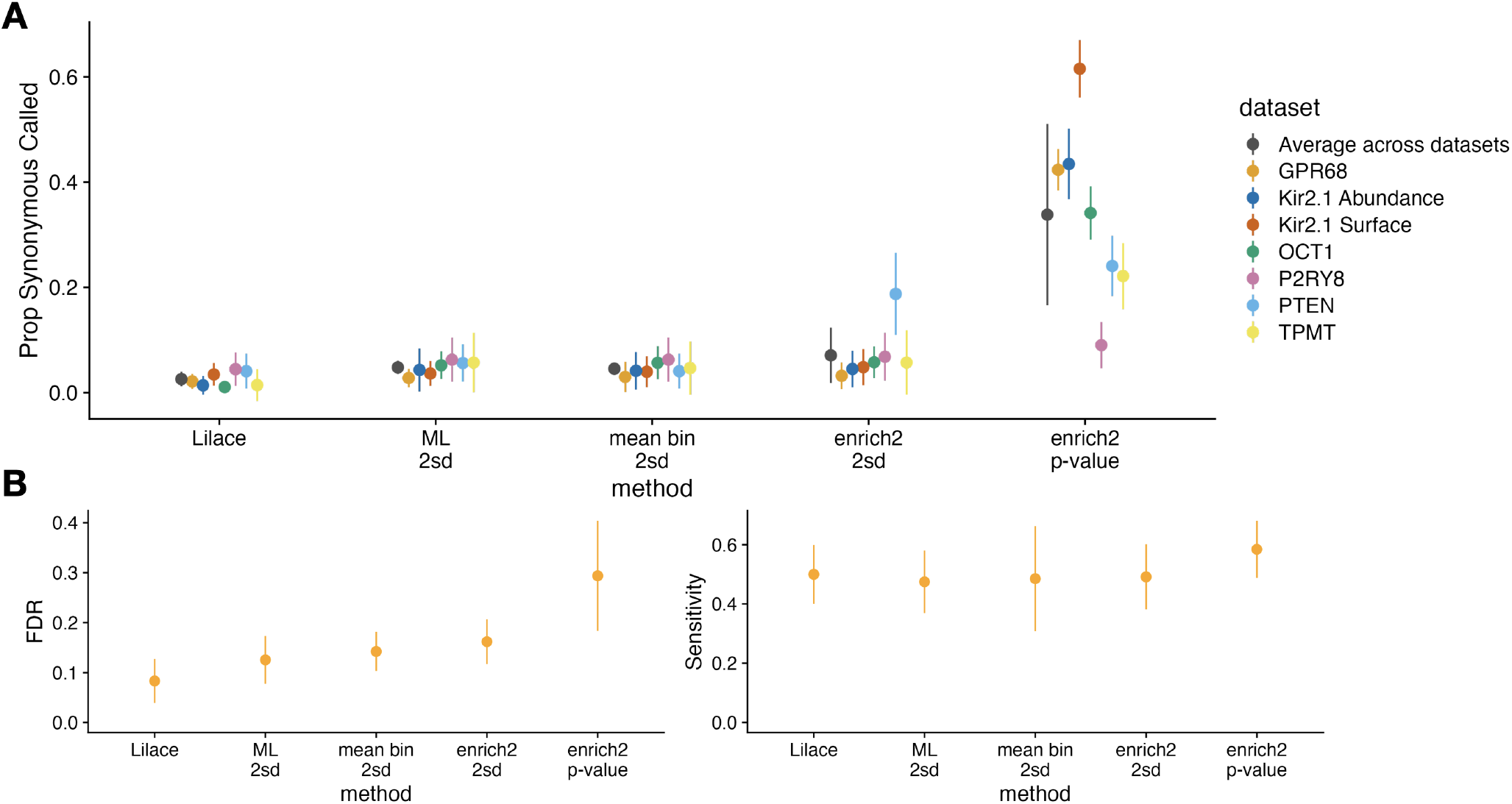
Empirical FDR and Sensitivity results for each discovery calling approach as determined by (A) masking 20% of the synonymous mutations to estimate FDR across 10 iterations and (B) using AlphaMissense pathogenicity labels across the first 6 datasets to determine relative FDR (benign label) and Sensitivity (pathogenic label). The P2RY8 dataset tracked poorly with AlphaMissense for all discovery calling approaches and is included separately in supplementary Fig. 9

### Lilace trims the discovery set in an OCT1 abundance screen

To qualitatively examine discovery differences, we compared Lilace’s results with those of previous approaches in the datasets used to seed our simulations. Lilace’s OCT1 abundance discovery set generally agreed with analysis from Yee et al. that used a two standard deviation cutoff of the Enrich2 scores, identifying the N-terminal transmembrane domains (TMs) as enriched for discoveries and several C-terminal TMs as depleted (Supp Fig. 11) (14). The discovery heatmap separates the regions enriched and depleted for effects (Fig. 5A, Supp Fig. 11), exemplifying how filtering by variant effect uncertainty can clarify biologically relevant patterns.

**Figure 5:**
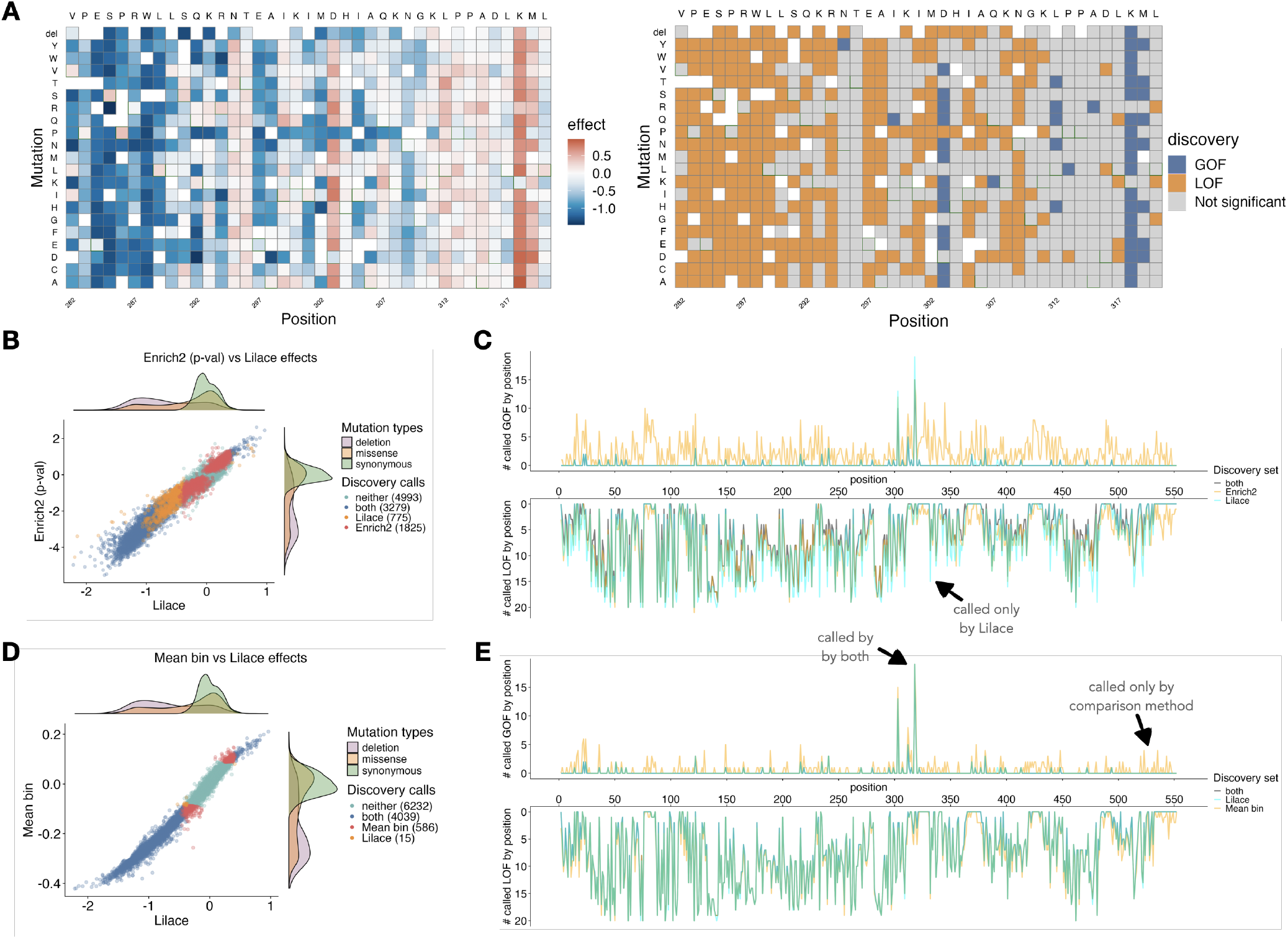
(A) Sample Lilace effect size and discovery heatmaps for OCT1 positions 282-320 at a significance threshold of lfsr † 0.05. In the discovery heatmap, orange colors indicate a significant LOF effect and blue colors indicate a significant GOF effect. (B) Score and discovery difference between Lilace and Enrich2 (p-value based), colored by whether a variant is discovered by both approaches, only Lilace, only Enrich2, or neither. (C) Position-wise discovery differences with Enrich2’s discovery set, comparing the number of significant GOF (upper) and LOF (lower) hits at each position. Orange indicates additional discoveries by the original approach, cyan indicates Lilace, and gray indicates both, with a resulting teal for overlapping discovery calls. (D-E) The same analysis as in B and C repeated for the weighted mean bin scoring approach.

We found that Lilace did not call around 13% (586/4625) of the discoveries called by the mean bin approach (Fig. 5D), which had the next best FDR in the simulations. These variants had sufficiently large variance in their effect sizes such that there was insufficient evidence for Lilace to call them as discoveries, reducing the overall discovery set. A stronger pattern in discovery differences emerged when compared with using the Enrich2 p-values, which is the only other approach tested that incorporates replicate variance and calls discoveries based on statistical significance. We expect Enrich2 to be misspecified for FACS data, as estimating a slope across bins does not take into account the shape of the underlying fluorescence distribution. This misspecification was reflected in the p-values, which only had a 0.68 correlation with the Lilace local false sign rates (lfsrs). In this case, we found that Lilace did not call around 36% (1825/5104) of the Enrich2-based discoveries (Fig. 5B). Additionally, we found Enrich2 had 24% (2600 / 10872) overall discovery call disagreement with Lilace, whereas the mean bin approach had 6% (601/10872) disagreement.

The reduction in called discoveries was especially noticeable in putative gain-of-function (GOF) variants, which are expected to be milder (31) and less frequent (1) than loss-of-function (LOF) effects. Lilace identified 94 total GOF effects, compared to 352 using the mean bin-based approach (Fig. 5E) and 1168 using Enrich2 (Fig. 5C), with substantial reduction in the terminal domains (Fig. 5C,E). Additionally, Lilace-identified GOF effects were largely concentrated at positions 303 and 318, of which the former was validated in follow-up experiments (14).

Lilace also identified an additional 15 and 775 variants compared to the mean bin approach (Fig. 5D) and Enrich2 (Fig. 5B), respectively. These were mainly LOF and scattered in positions with at least one other LOF substitution, demonstrating the effect of position-level shrinkage (Fig. 5C,E). We also found an additional GOF variant, I392F, which was not identified by either of the other two approaches and escaped negative position-level shrinkage (−0.54 position effect).

### Lilace trims discovery sets in different Kir2.1 phenotypes

To further examine Lilace’s generalizability and potential for discovery, we applied Lilace to the Kir2.1 dataset (12; 32). Whereas the OCT1 experiment was based on the VAMP-seq platform with fluorescent protein fusion to measure abundance, the Kir2.1 screen instead used an antibody fluorescent tagging protocol to measure surface expression and abundance (12). Since abundance is a precursor to surface expression, we expect there to be reasonable agreement between the two phenotypes, except for variants that affect only surface expression. The integration of discovery sets from both of these phenotypes illustrates the effect of Lilace’s discovery differences on downstream analyses such as phenotype comparison.

The Lilace discovery set highlighted the expected functional regions of Kir2.1, with much of the LOF signal concentrated in the pore domain and golgi export-related residues (Supp Fig. 13) (17). In both phenotypes, we found an overall reduced discovery set, consistent with our previous results showing Lilace’s prioritization of FDR control leads to a smaller discovery set. Compared with the mean bin scoring approach, Lilace had around 8% (894/11032) and 6% (646/11032) disagreement for the abundance and surface expression phenotype discovery calls, respectively (Fig. 6D, Supp Fig. Compared to the Enrich2 p-value-based discovery set, Lilace had a more substantial 36% (3953/11032) and 33% (3620/11032) discovery call disagreement for the two phenotypes (Fig. 6A, Supp Fig. 14). A substantial portion of the removed GOF discovery calls again came from the C-terminal domain

**Figure 6:**
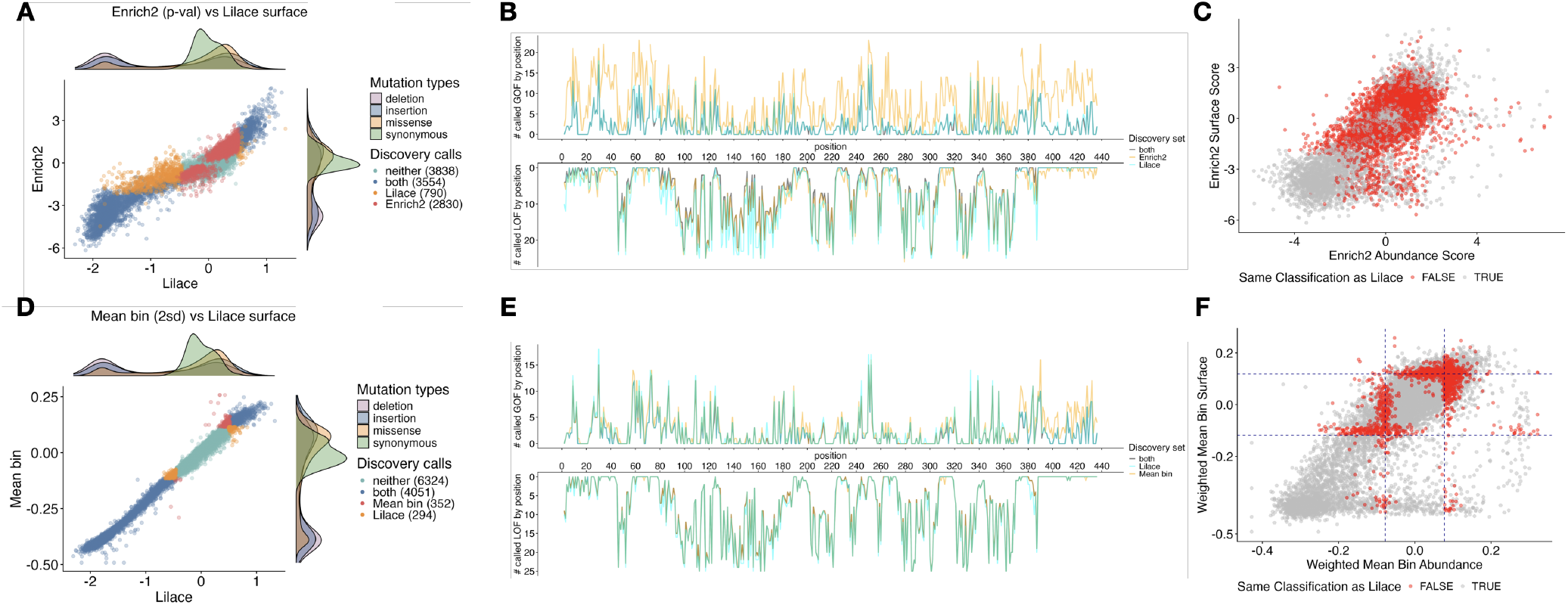
Lilace score and discovery classification differences with (A-C) Enrich2 p-value and (D-F) weighted mean bin for Kir2.1. (A,B) and (D,E) are Kir2.1 surface expression discovery comparison analogues to the previous OCT1 abundance plots. (C,F) compare the effects of both phenotypes on their original respective effect sizes, colored by disagreement with the Lilace discovery set. In (F), a two standard deviation cutoff on the original scores is plotted as a dashed line, showcasing the added flexibility of incorporating variant measurement uncertainty into the decision boundary.

(Fig. 6B,E), suggesting that the original calls could have been due to technical variance rather than biological signals. We observed greater measurement variance in this region, supporting this conclusion (Supp Fig. 15). While Lilace was more conservative in calling weaker-evidenced variants, it also discovered 377 additional variants across both phenotypes compared to the mean bin approach and 1431 additional variants compared to Enrich2. The most concentrated and consistent increase in discoveries occurred as additional LOF calls in the surface expression phenotype at positions 382-387 (Fig. 6E), which houses a previously identified diacidic ER-export motif (12; 17; 33).

The impact of a changed discovery set is compounded when model output is combined for downstream analysis, such as comparing phenotypes for mechanistic inference. When using our significance cut-off to classify variants as affecting only protein abundance, only surface expression, both, or neither, we found a 54% (5964/11032) overall difference with the Enrich2 p-value-based discovery set (Fig. 6C) and a 13% (1425/11032) overall difference with the mean bin-based set (Fig. 6F). Furthermore, compared to both methods, Lilace identified a lesser proportion affecting only abundance and a greater proportion impacting only surface expression or both phenotypes (Supp Table 2). These proportions align more with the phenotype hierarchy, since we expect variants that affect abundance to have a downstream effect on surface expression, but not the other way around.

## Discussion

Previously, no general statistical method had been designed to score FACS-based phenotypes, despite fluorescence being one of the most common readouts for DMS experiments. Furthermore, the performance of previous ad hoc scoring approaches had not been quantified, leaving their validity and generalizability across datasets unverified. As a result, variant scores from these experiments come without thorough quantification of their reliability and reproducibility, such as via standard errors and measures of significance like p-values. The lack of reliability and reproducibility can lead to inefficiency validating or interpreting variants with unknown score uncertainty. We fill this method gap with Lilace, a Bayesian hierarchical model that adapts its discovery classifications based on measurement uncertainty, improving on the reliability of previously applied ad hoc approaches.

Lilace leverages replicate variance and FACS-specific distributional assumptions for inference of variant effect size uncertainty, while using synonymous variant scores as an internal negative control set to mitigate false discoveries. Our results demonstrate that Lilace offers improved FDR control and robustness, based on both simulations and empirical benchmarks. In situations with increasing experimental variance, Lilace is the only approach to maintain a consistently low FDR at a given significance threshold. In situations with low experimental variance, the mean bin approach can also provide a reasonable discovery set, albeit without providing a clear cutoff for FDR control. On the other hand, Enrich2’s uncertainty quantification does not perform well in any scenario, which is consistent with the misalignment of growth-based modeling assumptions with FACS data.

By prioritizing FDR control, Lilace outputs a reduced discovery set in real data, especially in putative gain-of-function effects, likely due to fewer false positives and slightly lower sensitivity to small effects. While this mild decrease in sensitivity appears consistent, we expect that the large reduction in FDR will facilitate clearer interpretation of effects. Additionally, when the proportion of small effects is low, Lilace has no loss of sensitivity (Supp Fig. 3, 4, 5). Model extensions could help mitigate the sensitivity decrease with a less conservative inclusion of synonymous effect variance, such as an auxiliary model to learn the bias distribution from the negative controls (26). Improving synonymous variance incorporation could also improve the power gain from position-level shrinkage, since the shrinkage will apply to a greater proportion of the effect variance.

Like with Rosace, the position-level shrinkage boosts discovery power, without introducing substantial bias that prevents the identification of variant effects opposite to their position mean (22). Analogously to Rosace, this position grouping is only applicable to single or fixed mutation datasets, without straightforward application to random mutagenesis experiments with multiple mutations. While it is possible to run Lilace without the position-level grouping, properly extending Lilace to these datasets is an important area of future work. Although uncommon, a dataset may also not have a distribution of synonymous variants to use as a negative control. While it would still be possible to score against a given baseline proportion, such as a true wildtype or the overall fluorescence distribution, Lilace’s negative control-based bias correction is not possible and performance would likely echo that of unrecalibrated Lilace (Supp Fig. 10).

Recent work in DMS modeling has additionally incorporated priors on the effects of amino acid substitutions, which could be similarly incorporated into Lilace (34; 35). Other sources of prior information such as structure or protein language model embeddings could also be incorporated. Another potential extension is a variational approximation for computational efficiency (36). Finally, while we use cross-phenotype analysis to demonstrate the impact of improved discovery sets, a joint analysis framework is needed to fully model multi-phenotype relationships.

Although there is more work to be done, we hope that Lilace will enable researchers to obtain reliable, reproducible, and interpretable scores for FACS-based DMS experiments. This is especially critical as experiments scale to greater numbers of measurements across conditions, phenotypes, and proteins, making ad hoc analysis impractical. As we demonstrate in the multi-phenotype analysis, even seemingly small FDR differences can compound to change downstream interpretations of variant effects. We expect that more robust DMS analysis tools can aid functional dissection of genetic variant effects at every level, including protein mechanisms (1; 2), human phenotypes (5; 6), and clinical interpretation (7; 8; 9).

## Methods

### Data processing

Count datasets used for our analyses were received from their respective original publications (10; 12; 14; 18; 21). As a pre-processing step, we filtered out variants with less than 15 overall counts across replicates. We also added a pseudocount to help with model fitting.

In the scenario of unequal cell proportions in each bin, PCR amplification can lead to disproportionately large counts for bins with lower proportions. To account for this bias, we used the cell proportions from the FACS sort reports to normalize each bin’s counts to match its cell proportion (with the exception of PTEN and TPMT as sort reports were not available) (37). Given a variant *v* from replicate *r* with *K* bins, we rescaled the bin counts in bin *k*, *y*_*vrk*_, based on the replicate and bin-specific cell proportions π_*rk*_, the total read counts in that bin *R*_*rk*_, and the total read counts across bins *R*_*r*_.

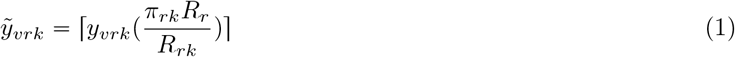

This results in a length *K* vector of rescaled bin counts for each variant and replicate: 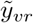. In the absence of cell sorting information for this normalization, Lilace’s use of a replicate-specific scoring baseline can still account for this bias (Supp Fig. 18).

### Model details

Lilace models the bin counts 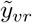 from variant *v* and replicate *r* as,

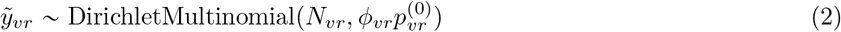

where *N*_*vr*_ is the sort proportion-normalized variant read count, *ϕ*_*vr*_ is the overdispersion term, and 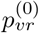 is the variant’s latent bin probability across *K* bins. We model *ϕ*_*vr*_ using a linear trend function of the variant counts, estimated within each replicate,

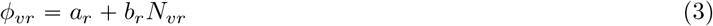

To estimate an effect size for each variant from *K* gating bins, we model the replicate-specific baseline fluorescence bin probability of the negative controls (synonymous variants) *q*_*r*_ and estimate the distance of each 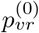 from *q*_*r*_ as a shift in the corresponding probability quantiles of a standard normal,

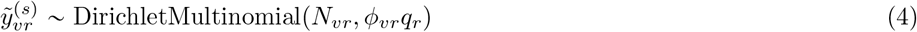

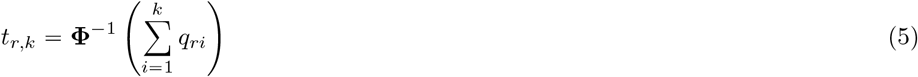

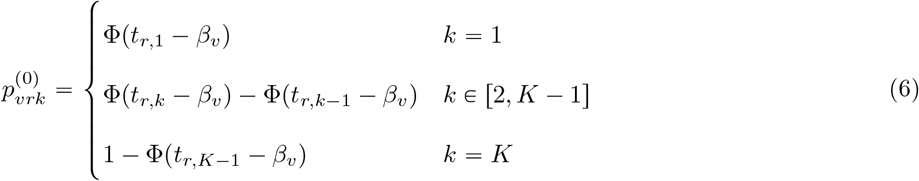

where *y*^(*s*)^ is a subset of *y* containing only the synonymous control variants, *t*_*r,k*_ is the standard normal quantile cutoff corresponding to bin *k*’s baseline probability in replicate *r* for *k* ∈ [1,K− 1], and Φ (·) is the standard normal CDF. The effect size (*β*_*v*_ is the shift in these cutoffs to match variant v’s bin probabilities. Similar latent normal quantile approaches have previously been successfully applied to FACS-based CRISPR screens (23; 38; 39). We jointly estimate *q*_*r*_ and *β*_*v*_ during estimation to capture variance in the baseline cutoffs. We find that modeling a replicate-specific baseline instead of a global *q* can account for bias due to replicate-specific shifts in the counts, such as due to unequal sorting or PCR amplification (Supp Fig. 18).

We incorporate position level hierarchy by sampling effects from a position-specific distribution:

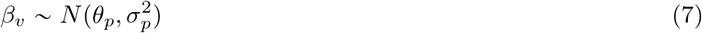

where *θ*_*p*_ and 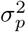 represent the mean and variance of the effects at position *p*. This parameterization results in position-specific shrinkage, meaning effects incorporate a prior bias towards their position’s mean effect to help estimation in low sample size variants. While effect shrinkage is helpful for estimating variant true effects, we aim to avoid excessive shrinkage in synonymous variants to preserve their interpretation as a negative control group. To this end, Lilace encodes a fixed effect for each synonymous mutation, so that each has their own mean, while still sharing a single variance term.

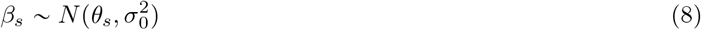

where s indexes the set of synonymous mutations. We then use the individual posteriors of each synonymous mutation for negative control-based bias correction on our effect sizes (26). We incorporate the variance between synonymous mutations into our decision boundary to separate variants into those that act as wild type-like and those with a substantial impact. Without this effect recalibration, we find that the model does not have adequate FDR control (Supp Fig. 10). Let *D* be the number of posterior samples and *S* be the number of synonymous variants. For each posterior sample 1,…, *D* and variant *v*, we select a random synonymous variant *s*_*vd*_ to recalibrate that posterior sample’s score.

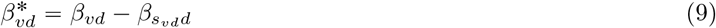

Our new score is 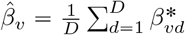 with variance 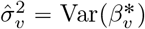 To classify an effect as a discovery we use the local false sign rate (lfsr) of the corrected posterior 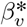 (40), with a default discovery threshold of 0.05.

The position and synonymous mean parameters *θ*_*p*_ and *θ*_*s*_ are given *N*(0, 1) priors. The variance parameters σ_*p*_ and σ_0_ are given InvGamma(1, 1) priors. The baseline threshold probabilities *q*_*r*_ are given a Unif(0, 1) prior. All other parameters are given a flat uniform prior. We fit Lilace with MCMC sampling using the default NUTS algorithm for HMC (41) in Stan (42). On most proteins, Lilace runs within a couple of hours (Supp Fig. 19).

### Other method details

We ran Enrich2 with each bin encoded as a timepoint and using the sum of synonymous variant counts in each bin as the wildtype reference. We used either the provided p-value with a Benjamini-Hochberg FDR correction (43) or standard deviations from the synonymous score as the discovery threshold.

For the weighted mean bin approach, we computed the score for a variant as:

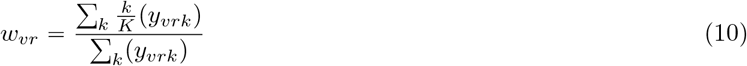

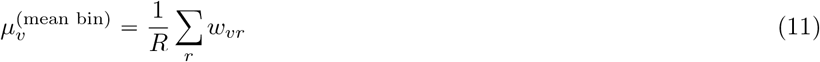

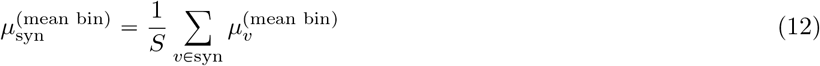

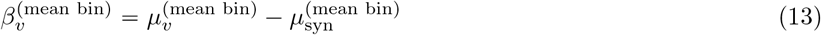

For the maximum-likelihood approach, we estimated the underlying fluorescence distribution using the raw FACS gating thresholds by minimizing the negative log-likelihood of a log-normal (28).

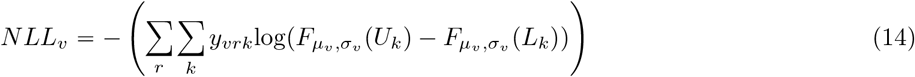

where *F* is the cumulative distribution function for a log-normal, *U*_*K*_ and *L*_*k*_ represent the upper and lower FACS gating thresholds for bin *k*, and *µ*_*v*_ and σ_*v*_ represent the mean and variance parameters of the log-normal.

We then computed a score as the difference in means between the average mean fluorescence of a variant and that of synonymous variants:

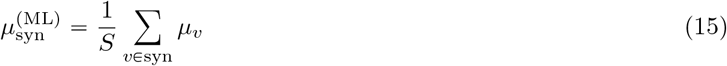

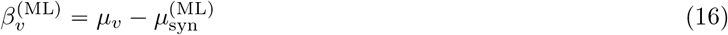

For both the weighted mean bin and maximum-likelihood based approaches, we set a discovery threshold using the synonymous variant score distribution, equivalent to a fixed standard deviation-based cutoff. This approach performed better than approximating the standard error of (*β*^(mean bin)^ or (*β*^(ML)^ with a normal distribution (Supp Fig. 10). To compute the discovery threshold, we calculated a p-value for the score by fitting a normal distribution to synonymous variant scores as a null distribution. For example, a 2 standard deviation cutoff of the synonymous distribution corresponds to approximately a 95% confidence interval or a p-value cutoff of around 0.05. We did not apply a multiple testing correction, to align with the way the standard deviation cutoff is commonly employed.

### Simulation methods

To compare and benchmark Lilace with simulated data, we designed a separate generative model of the experimental process that parameterizes various sources of natural and technical variance. Since this model was used only for simulation purposes, we did not need to ensure identifiability and implemented a more parameterized model. We marginally estimated simulation model parameters from real data to provide more realistic defaults, while toggling each one across a range of values to examine the relative performance of our model with other approaches. We determined discoveries based on a 95% certainty threshold for all approaches (0.05 p-value threshold for frequentist methods).

Our simulation procedure was as follows:

1. Input protein and experimental summary statistics
2. Simulate ground truth effect sizes
3. Simulate variant cell fluorescence and bin read counts
4. Add experimental variance in observed counts

We first picked an input protein dataset and set default simulation constants based on this protein including *V* variants per position, *R* replicates, *L* total read counts, and *K* FACS bins. We also set the default number of cells per variant, *m*, to 200. For simulation efficiency, we fixed the number of positions *P* to 100.

To estimate the effect size distribution from data, we applied the Log Normal MLE approach, since these effect sizes are on the same scale as the underlying fluorescence distribution’s parameters. However, since the individual variant effect sizes were unreliable due to low sample size, we only used the summary statistics of the collective effect distribution to parameterize our simulation. To enable the investigation of different effect size architectures via toggling the proportions of small effects, we estimated separate distributions for small and large effects. We first fit a bivariate Gaussian mixture model to the maximum-likelihood derived effect sizes to estimate a distribution of small effects and a distribution of large effects, labeling the small effect mode as the one containing more synonymous variants. From the fitted mixture model, we derived the proportion of small effects *p*_small_ and fit a Gaussian to each mode to derive the parameters of both effect groups, *µ*_small_, *σ*_small_, *µ*_large_, *σ*_large_. Note that with this procedure many of the small effects are derived from variants that are unlikely to have a significant effect in the real data, but incorporating simulated effects of such a low magnitude helps compare scoring approaches in benchmarking. Next, we simulated from these fitted groups with (*β*_small_ ∼ *N* (*µ*_small_, *σ*_small_) and (*β*_large_ ∼ *N*(*µ*_large_, *σ*_large_). We truncated the effects to be within 2 standard deviations of their group mean to remove outliers and create a greater distinction between the groups.

We then used these effect sizes to generate ground truth effect sizes in a position-specific manner. We parameterized the position effect as the variance in effects at a position σ_*p*_, which we estimated from the MLE-based effect sizes. At each position *p*, we decided the effect pattern based on the real data’s effect pattern *W* determined using the weighted mean bin approach. Every synonymous variant or variant that is not identified as an effect in the real data was assigned a zero effect size. We sampled the rest of the effects by first uniformly sampling a position mean *θ*_*p*_ from our effect size distribution, with a probability *p*_small_ of sampling from (*β*_small_ and a probability 1 − *p*_small_ of sampling from (*β*_large_. Then, we sampled the effects at that position as (*β*_*p*_ ∼ *N*(*θ*_*p*_, σ_*p*_). Under large σ_*p*_, the proportion of small and large effects at the position could change; however, on average these proportions stayed consistent across the simulated dataset.

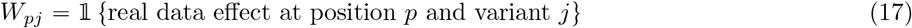

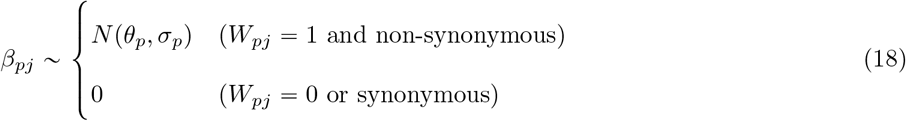

To optionally toggle the overall proportion of effects, we took *W* and randomly increased or decreased the effects until the target effect proportion was reached.

To simulate each variant’s latent fluorescence distribution, we first used the set of synonymous variants to estimate the fluorescence distribution of wild type-like variants using the Log-Normal MLE approach, treating each synonymous count observation as an independent observation of the same distribution. Let *µ*_0_ and *τ* be the wild type-like Log-Normal location and scale parameters. For each simulated variant *v*, we sampled m cells from the latent fluorescence observation *F*_*v*_ as:

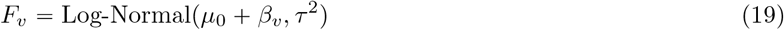

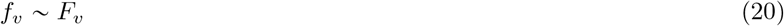

When simulating a replicate effect *B*, we added a constant term *b*_*r*_, derived from length *R* equidistant steps in [− *B, B*], to the log-normal location parameter in replicate *r*. We then set gating thresholds as equally spaced quantiles of the overall fluorescence distribution Σ_*v*_ *f*_*v*_ and divided cell counts into sorting bins accordingly. We used those latent sorted cell counts to set the true bin probability for a variant 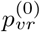.

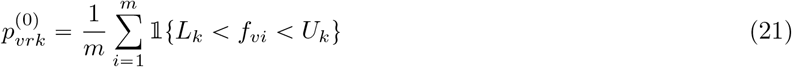

where *L*_*k*_ and *U*_*k*_ are the lower and upper fluorescence bounds of bin *k*, respectively. When simulating shifted gates, we shift non-endpoint *L*_*k*_ and *U*_*k*_ by a constant gate offset term on a log scale (to match the log normal effect magnitudes).

To generate the observed counts, we incorporated variant-specific technical variance by sampling from a Dirichlet-Multinomial with overdispersion parameter *η*_*v*_. Since only a handful of replicates is insufficient to estimate the parameters of a Dirichlet-Multinomial, we used the set of synonymous variants to estimate the relationship between variant read count and *η*_*v*_. We first estimated the expected synonymous bin probabilities

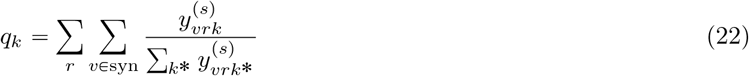

To get estimates of *η*_*v*_, we used Brent’s method with a lower bound of zero and an upper bound of 1000 to minimize the negative log-likelihood:

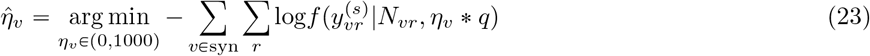

where *f* is the Dirichlet-multinomial likelihood and the bin proportions *q* are fixed to their expected value. We then regressed these estimates of *η*_*v*_ on the average replicate read count for that variant 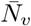 to get a linear estimate of the count-variance function.

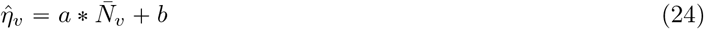

For each replicate, we then randomly sampled a read count 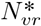 from the seed dataset, plugged 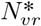 into equation to estimate 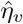, and simulated variant *v*’s observed bin read counts as

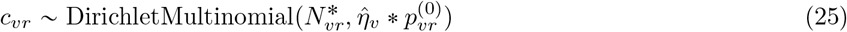

where *c*_*vr*_ is variant *v*’s cell count in replicate *r* across *K* bins.

When benchmarking performance across bin overdispersion parameters, e.g. due to PCR amplification bias, we instead set *c*_*vr*_ to the simulated sorted cell counts, then sample the reads from bin *k* with an input overdispersion parameter *ψ*_*k*_:

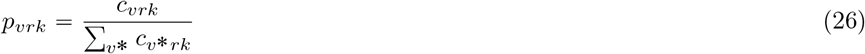

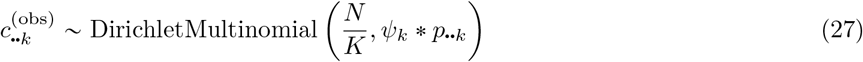

We used the resulting counts to benchmark our model’s performance and make relative comparisons with other approaches. When toggling specific parameters, we set that parameter to a specific parameter and left the others at their data-estimated defaults. Each simulation setting was repeated 10 times to obtain average FDR and sensitivity statistics with standard errors.

### Empirical Benchmarking

For our synonymous-based empirical benchmarking (Fig. 4A), we randomly masked 20% of synonymous mutations in each dataset tested, treating them as any other variant at their position. We estimated FDR by computing the fraction of masked synonymous variants called and ran this procedure 10 times with different masks to get standard errors. For the AlphaMissense-based benchmarking (Fig. 4B), we retrieved available AlphaMissense pathogenicity predictions and scores for each protein tested (44), then used benign labels as a proxy true negative and pathogenic labels as a proxy true positive to compute FDR and sensitivity on. For the ClinVar-based sensitivity benchmarking of Kir2.1 and PTEN (Supp Fig. 8), we used missense variants labeled “likely pathogenic” or “pathogenic” as our proxy true positive set.

## Declarations

### Ethics approval and consent to participate

Not applicable.

### Consent for publication

Not applicable.

### Availability of data and materials

Lilace is available to run as an R package and can be installed from https://github.com/pimentellab/lilace. Retrieved count data and code used for simulations and analyses can be found at https://github.com/jermoef/lilace-paper-analysis. Original datasets for OCT1 (14), Kir2.1 (12), GPR68 (21), PTEN (10), TPMT (10), and P2RY8 (18) can be retrieved from their respective publications. ClinVar pathogenicity labels (30) (https://www.ncbi.nlm.nih.gov/clinvar/) and AlphaMissense scores (44) (https://alphamissense.hegelab.org/) were obtained from their respective online resources.

### Authors’ contributions

JF, JR, WCM, and HP conceived the project. JF, JR, and HP developed the statistical model. JF and HP designed the simulations. JF performed benchmarking and data analysis, with input from JR, MH, CM, NG, WCM, and HP. JF and HP wrote the manuscript, with input from JR, MH, CM, NG, WCM, and HP. All authors read and approved the final manuscript.

## Acknowledgements

The authors thank the HP and WCM labs for their many stimulating discussions. This work used computational and storage services associated with the Hoffman2 Cluster which is operated by the UCLA Office of Advanced Research Computing’s Research Technology Group.

