## Supplementary Figures for "Accurate variant effect estimation in FACS-based deep mutational scanning data with Lilace"

### Supplement

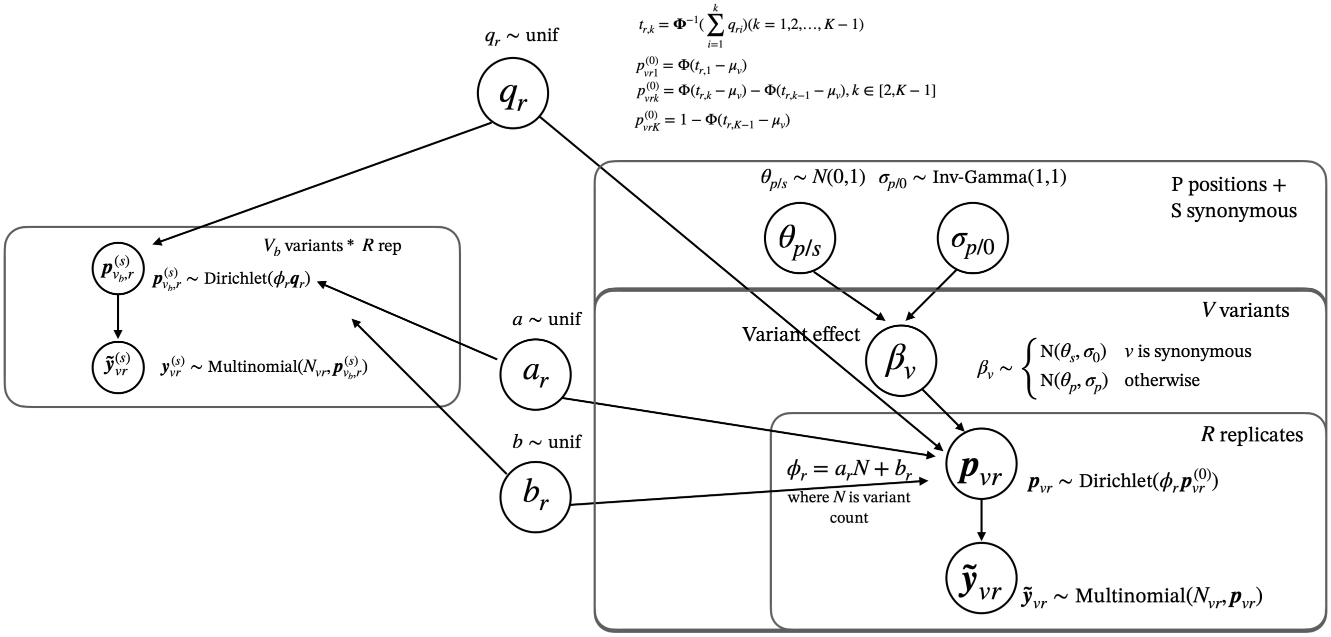


Figure 1: Bayesian plate model for Lilace

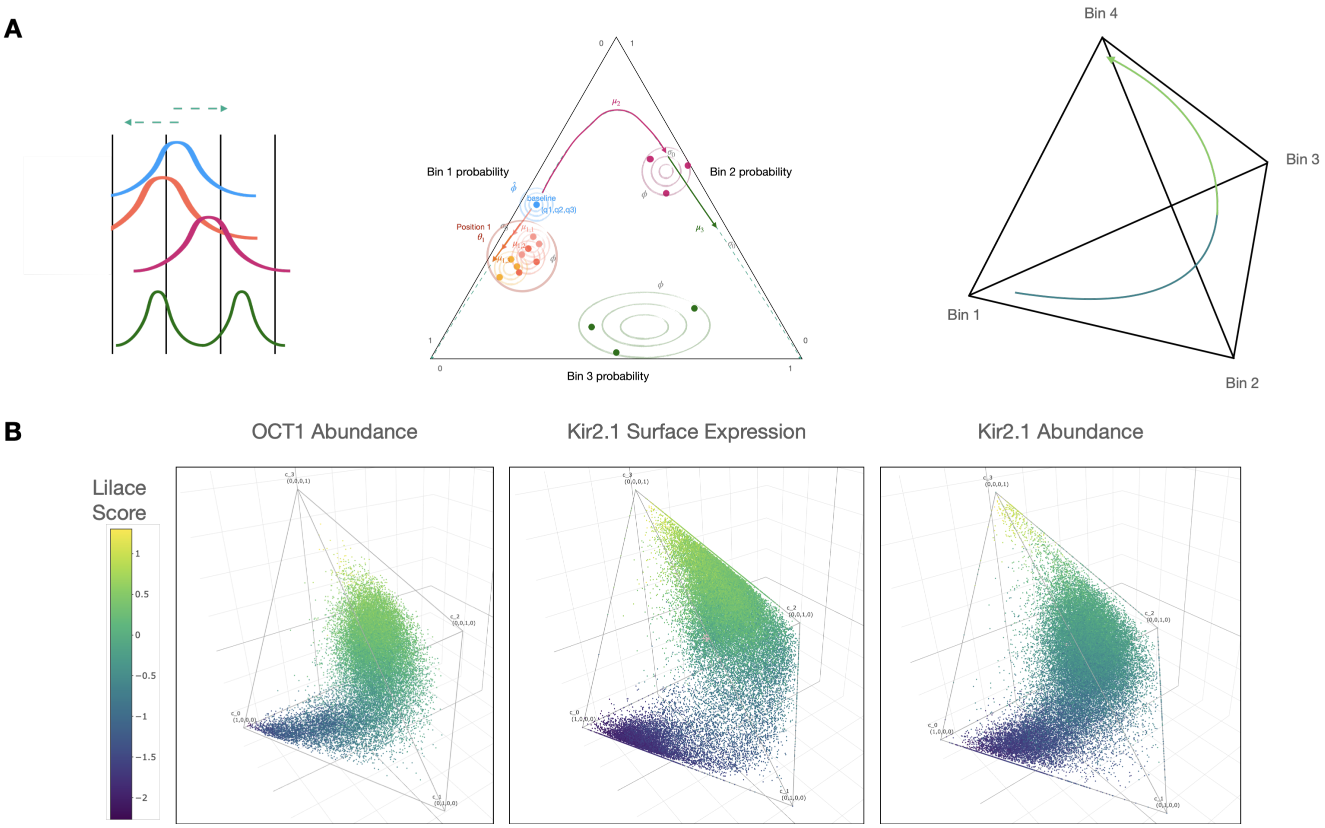


Figure 2: (A) Visual representation of latent effect size trajectory in probability space for $K = 3$ or $K = 4$. (B) Plots of the raw count data proportions in the same space depict a similar trajectory across variants. Each variant observation is a point that is colored by its score, with low scores having a large count proportion in the lowest bin ($c_0$) and high scores in the highest bin ($c_4$). A bimodal variant would be opposite of the curve (most counts spread across the first and last bin)–we can see there are very few, if any, such variants, and they are assigned a moderate score.

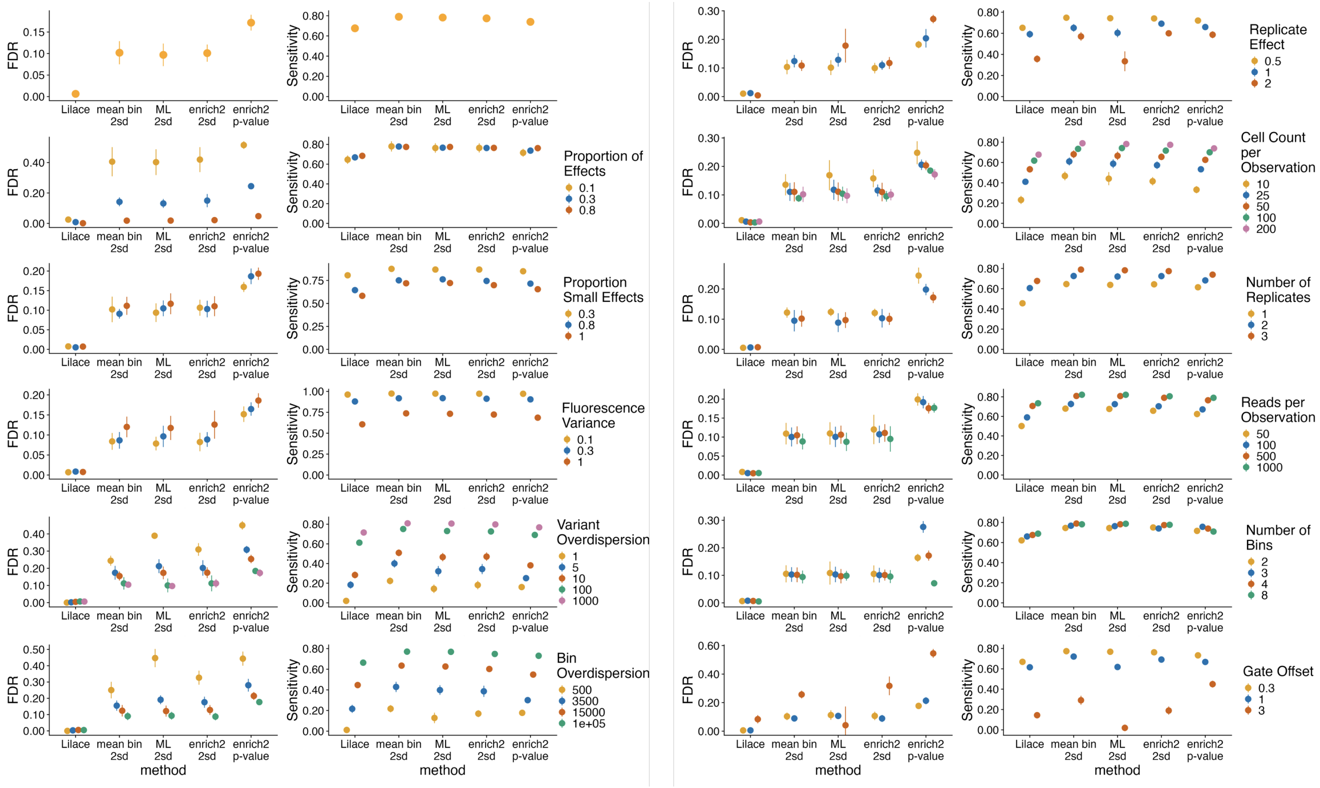


Figure 3: Full OCT1 Abundance Simulations

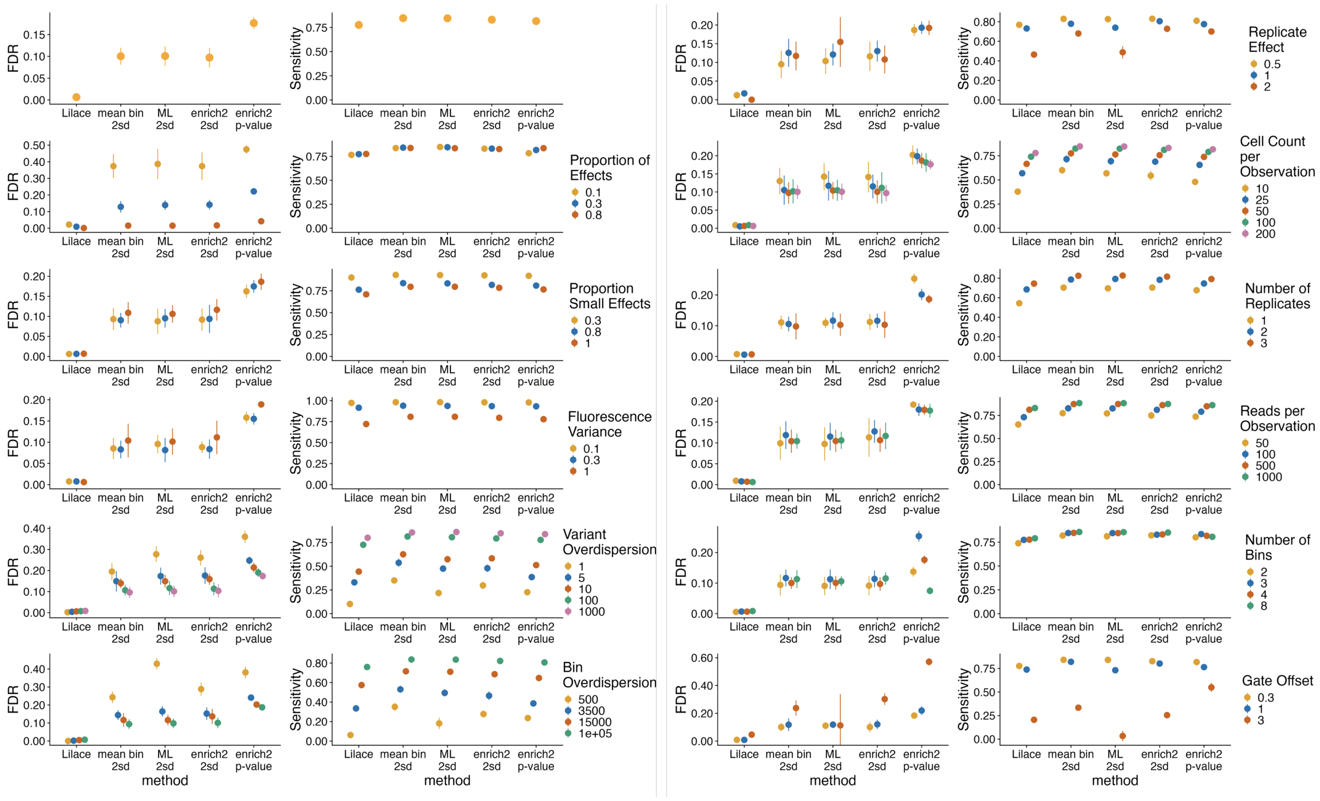


Figure 4: Full Kir2.1 Surface Expression simulations

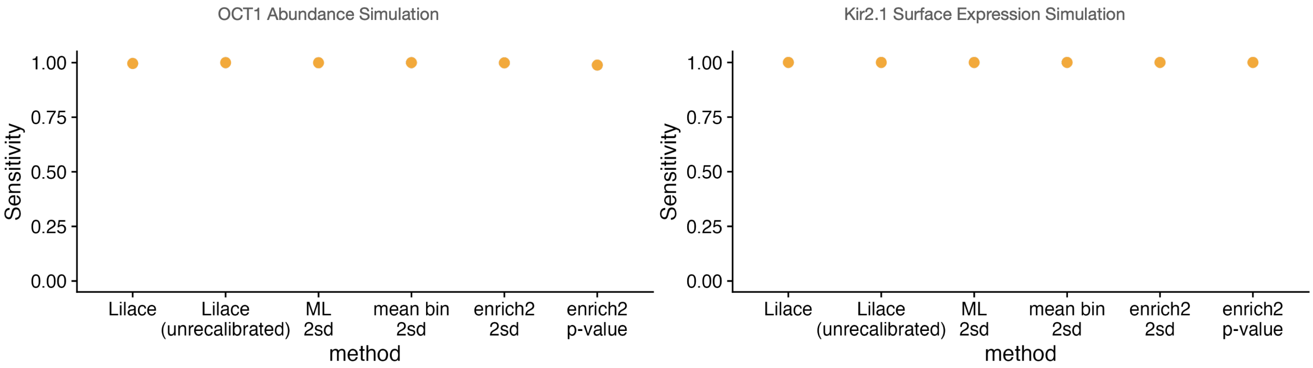


Figure 5: Default simulation sensitivity results filtered to the top 50% of effects

| Dataset | Lilace Correlation | Mean bin correlation | ML correlation | Enrich2 correlation |
|---|---|---|---|---|
| OCT1 | **-0.61** | **-0.61** | -0.59 | -0.59 |
| Kir2.1 surface | **-0.38** | **-0.38** | -0.37 | -0.34 |
| Kir2.1 abundance | **-0.36** | -0.35 | -0.35 | -0.31 |
| GPR68 | **-0.70** | -0.63 | -0.59 | -0.57 |
| PTEN | **-0.44** | -0.34 | -0.40 | -0.35 |
| TPMT | **-0.62** | **-0.62** | -0.59 | -0.59 |
| P2RY8 | -0.59 | -0.59 | -0.59 | -0.59 |

Table 1: Scoring approach correlations with AlphaMissense pathogenicity scores (higher AlphaMissense scores indicate higher pathogenicity probability)

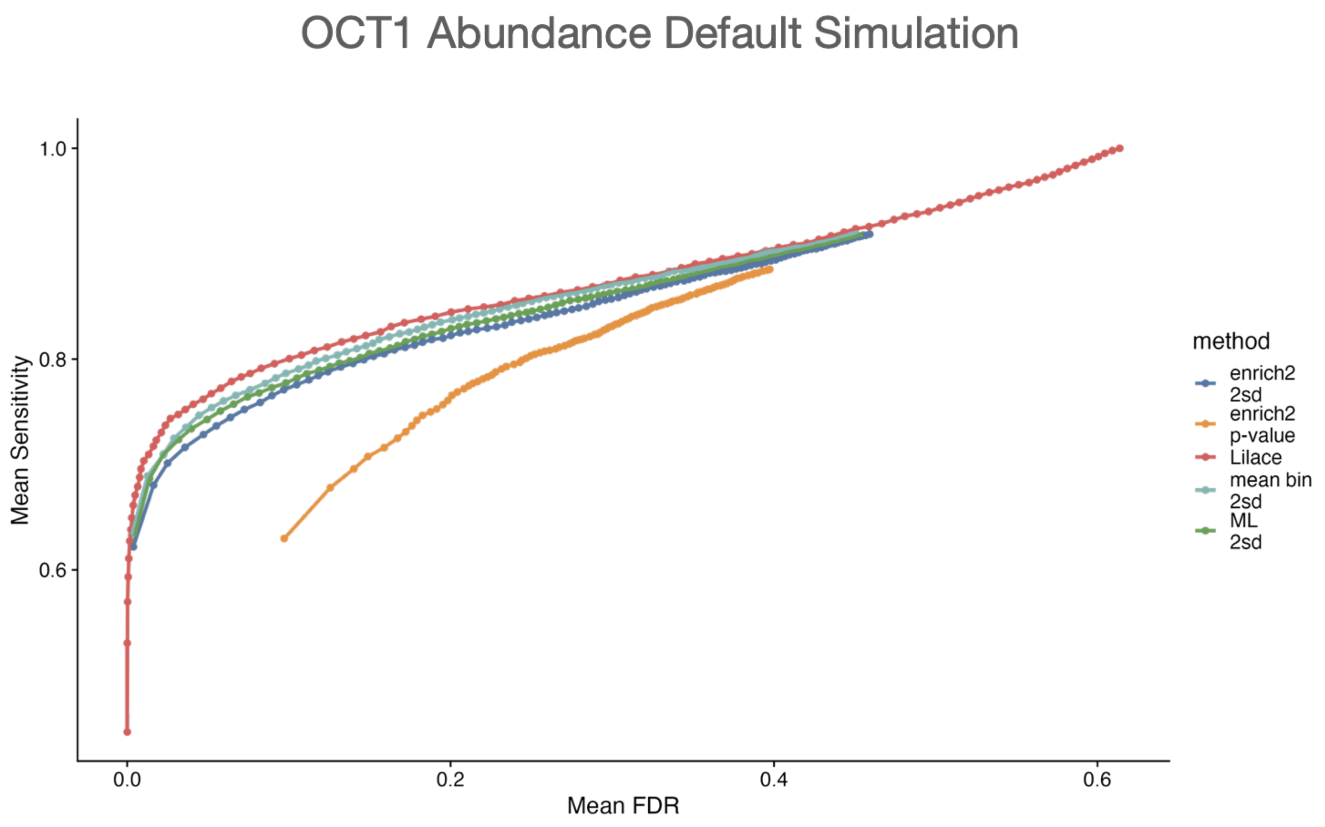


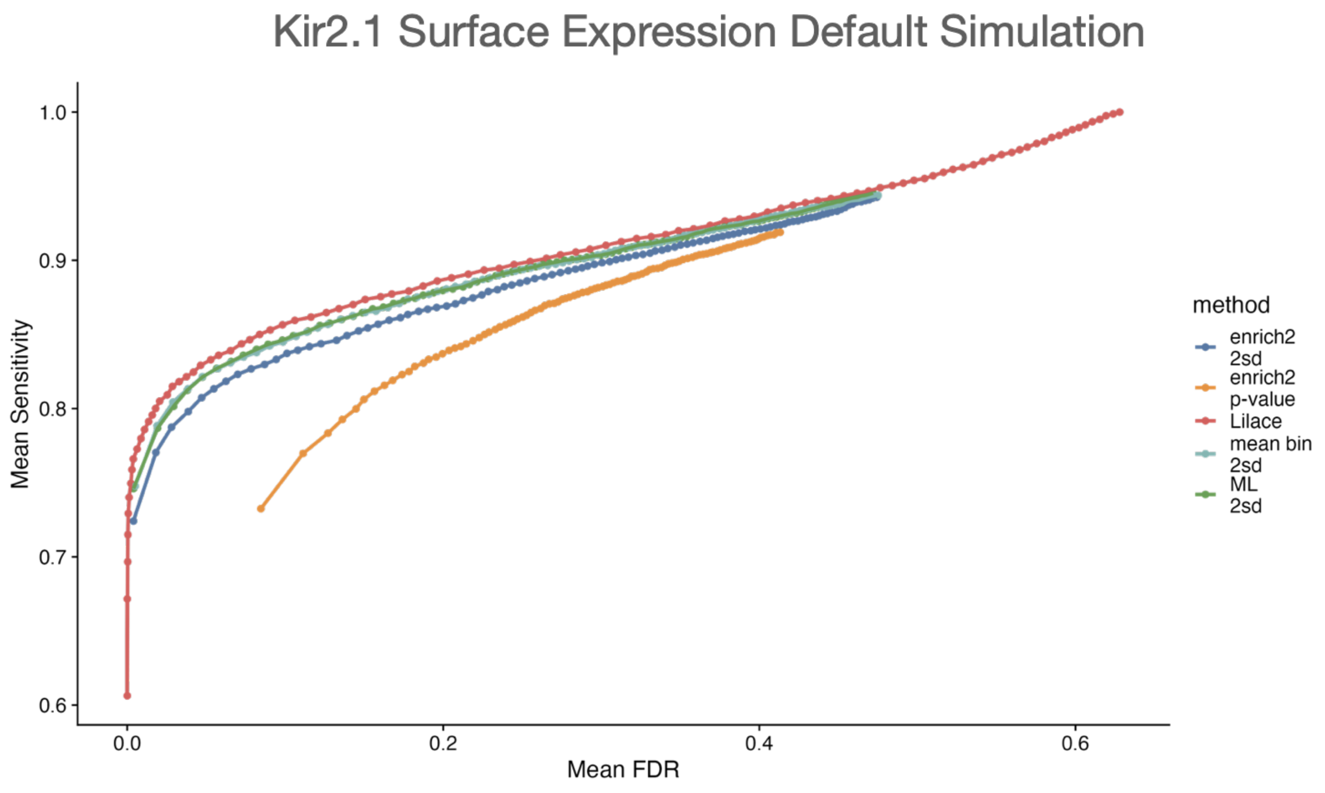


Figure 6: FDR-sensitivity tradeoff curve that plots the sensitivity for a given FDR level for each approach in the default simulation setting. Each point represents a significance threshold ranging from 0.001 to 0.5. Lilace also edges out other approaches by this metric, achieving the highest sensitivity for a given FDR level.

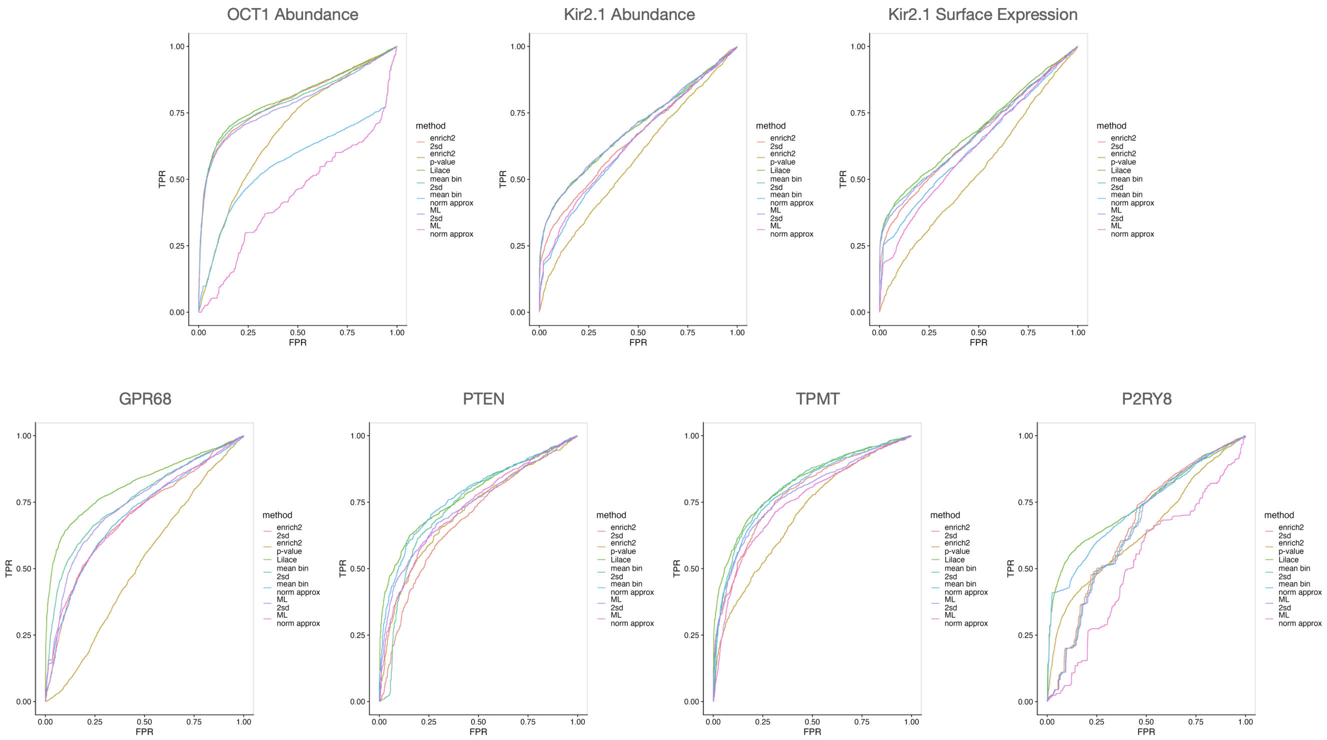


Figure 7: ROC curves for each approach based on AlphaMissense pathogenicity levels. Lilace is the only method that is consistently a top performer in each dataset.

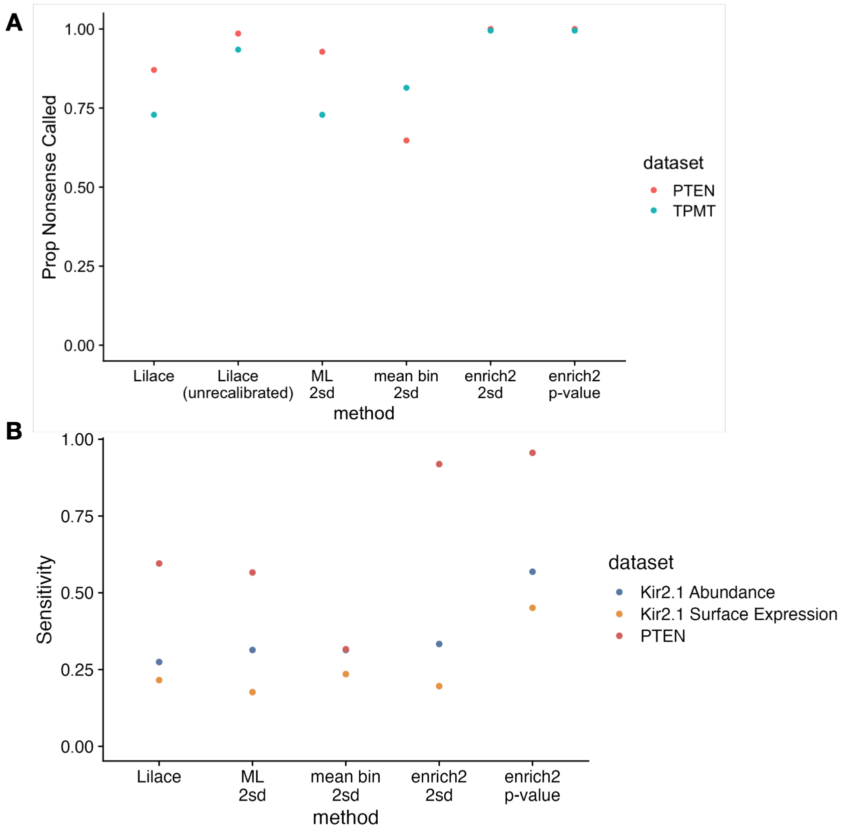


| Protein | # Benign Missense | # Pathogenic Missense |
|---|---|---|
| OCT1 | 6 | 0 |
| Kir2.1 | 2 | 51 |
| GPR68 | 3 | 1 |
| PTEN | 5 | 224 |
| TPMT | 3 | 0 |
| P2RY8 | 0 | 1 |

Figure 8: (A) Nonsense and (B) pathogenic ClinVar sensitivity checks when available. Only Kir2.1 and PTEN had enough labeled ClinVar variants to be used for this metric.

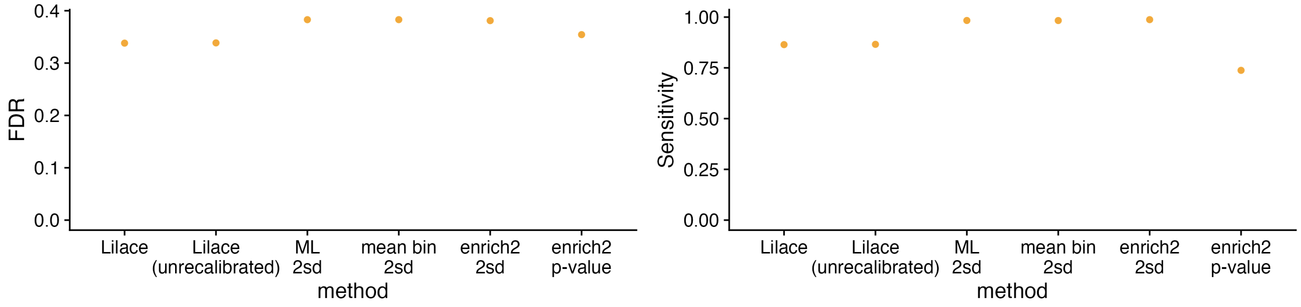


Figure 9: P2RY8 AlphaMissense-based FDR and sensitivity comparisons

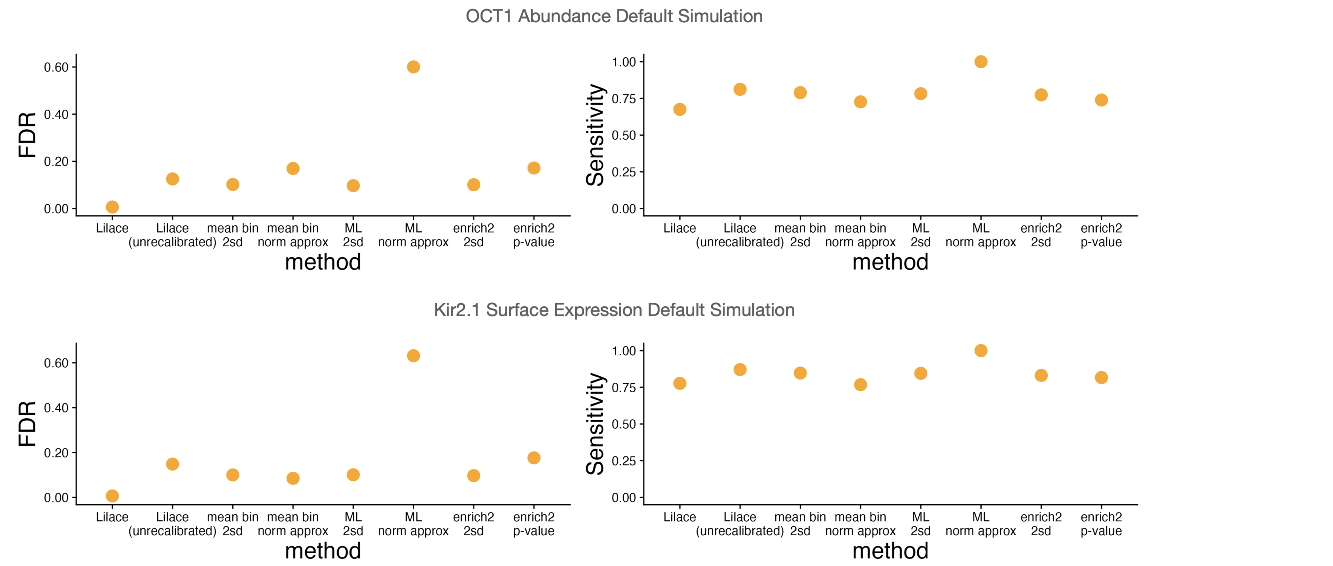


Figure 10: Full method comparison set in the default simulation setting, including unrecalibrated Lilace and normal approximations based on the score standard deviation for the ML and mean bin approaches.

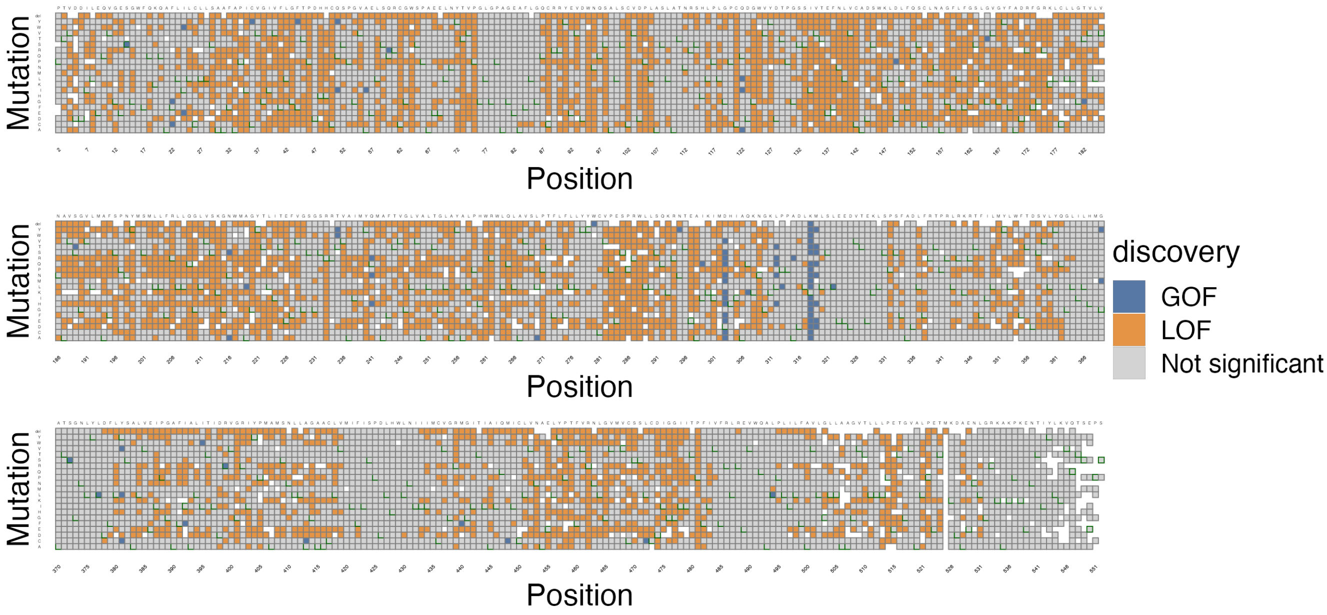


Figure 11: OCT1 full Lilace discovery heatmap

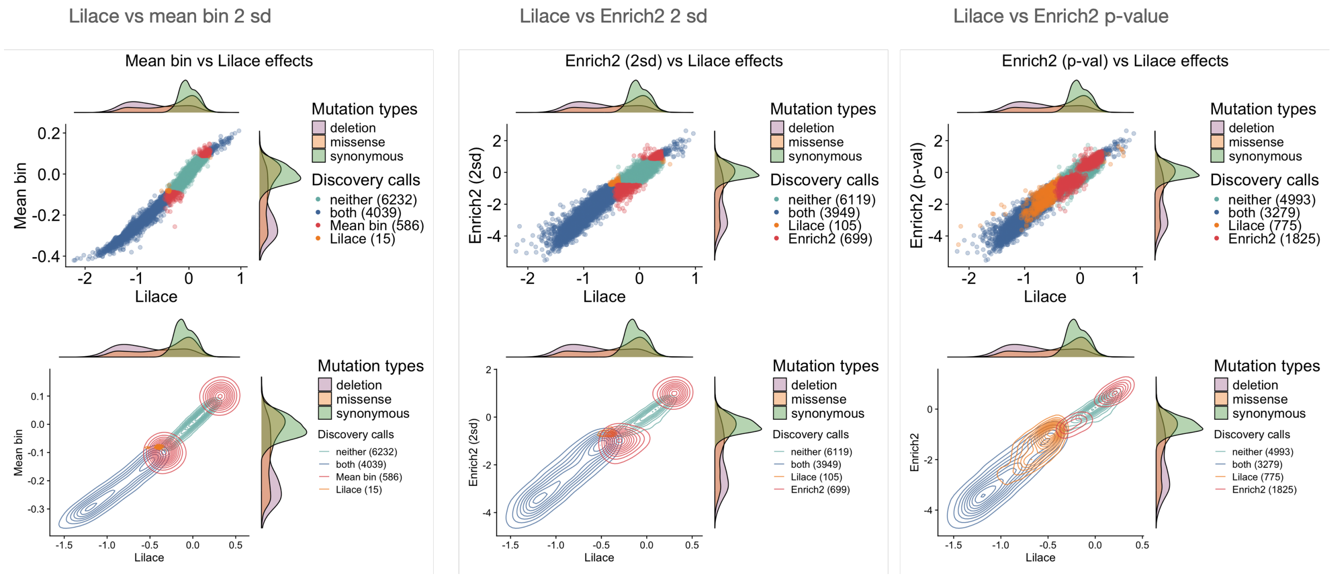


Figure 12: OCT1 real data method comparisons

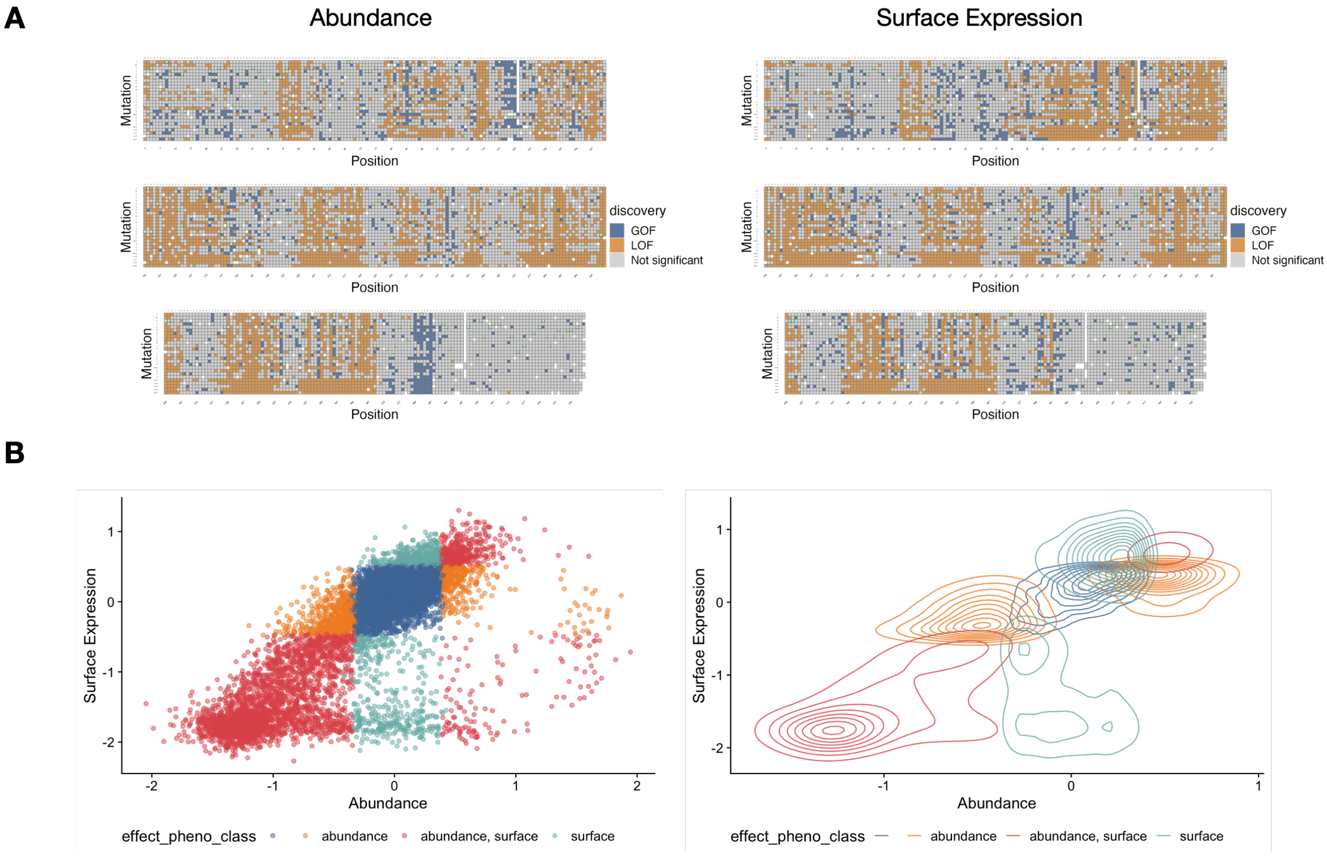


Figure 13: Kir2.1 Lilace discovery heatmaps and phenotype comparison plots

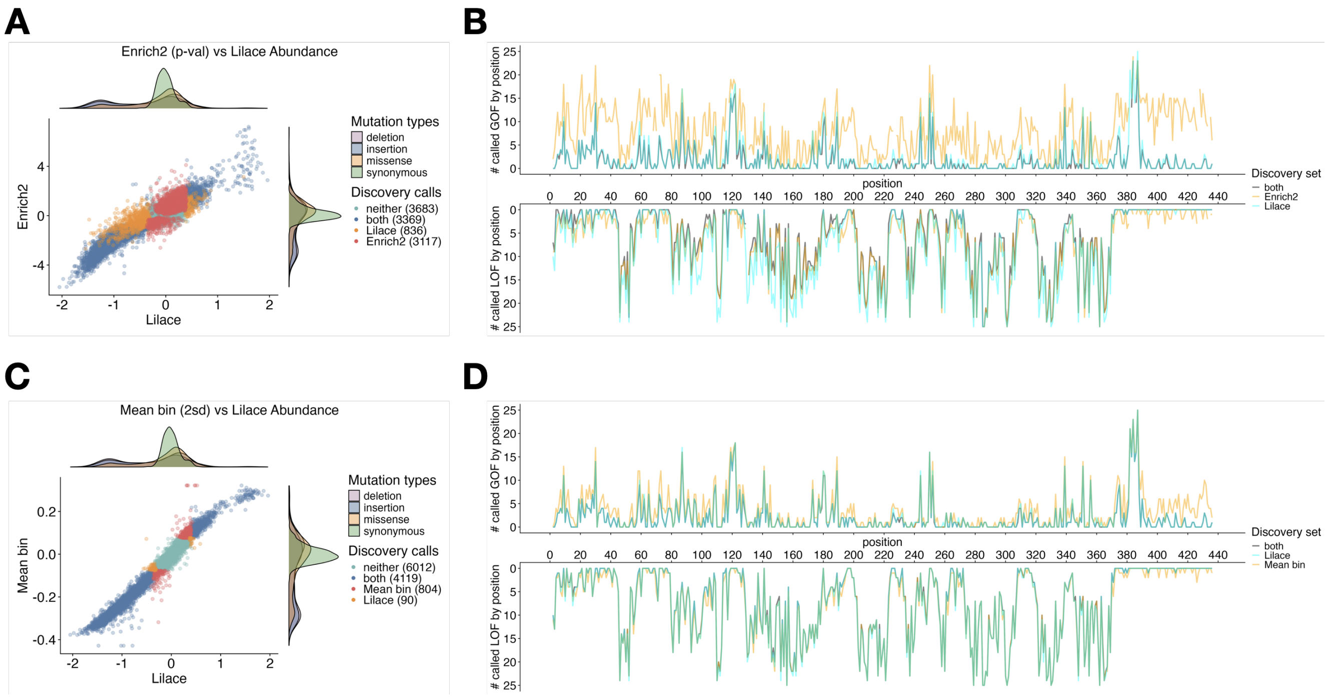


Figure 14: Kir2.1 Abundance method comparison

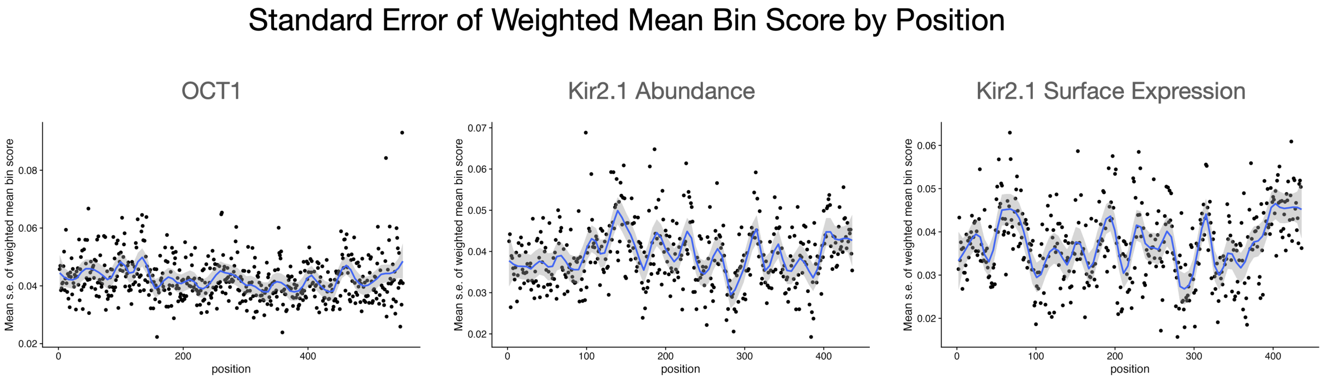


Figure 15: Average standard errors of Weighted Mean Bin Score by position. Standard error is computed as standard deviation of the score across the three replicates. Smoothed line is fit with a loess curve with a span of 0.1.

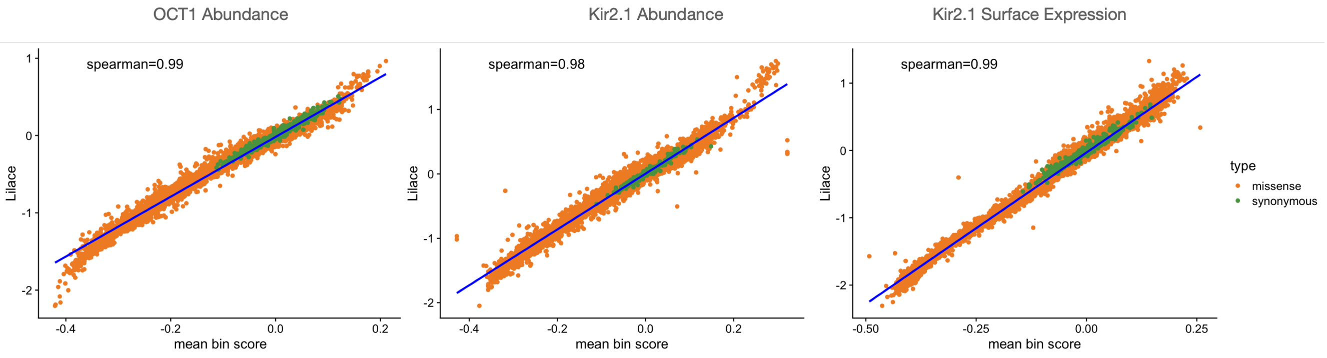


Figure 16: Bin rank consistency between Lilace and the weighted mean bin approach

| Method | Abundance Only | Surface Expression Only | Both |
| --- | --- | --- | --- |
| Lilace | 908 | 1042 | 3299 |
| Enrich2 p-value | 2165 | 2059 | 4317 |
| Mean Bin 2sd | 1468 | 948 | 3448 |

Table 2: Kir2.1 joint phenotype discovery numbers

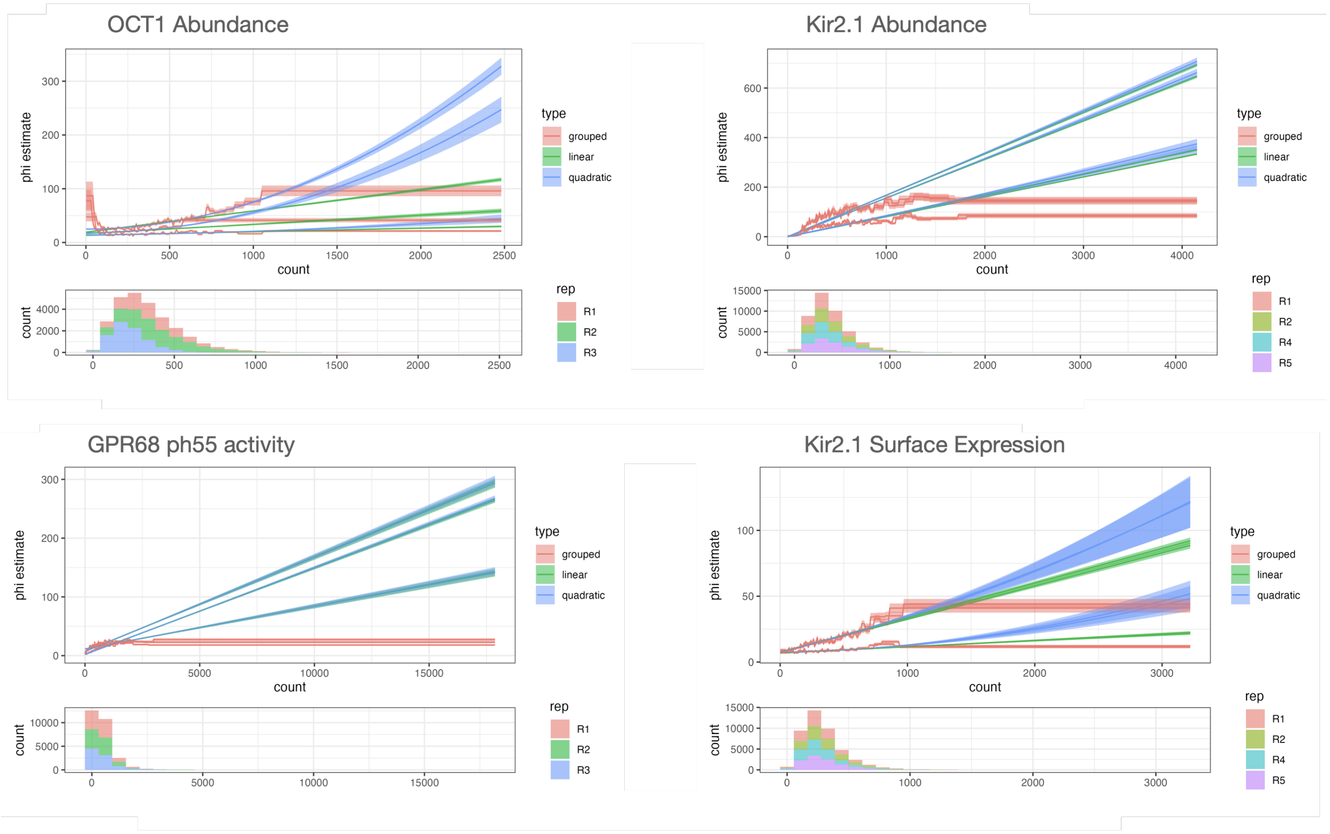


Figure 17: Comparison of count-overdispersion trend function estimates. In the regime of counts where almost all variant observations are located, the linear function approximation is similar to a employing a quadratic approximation or a Rosace-like non-parametric count grouping of size 100 each.

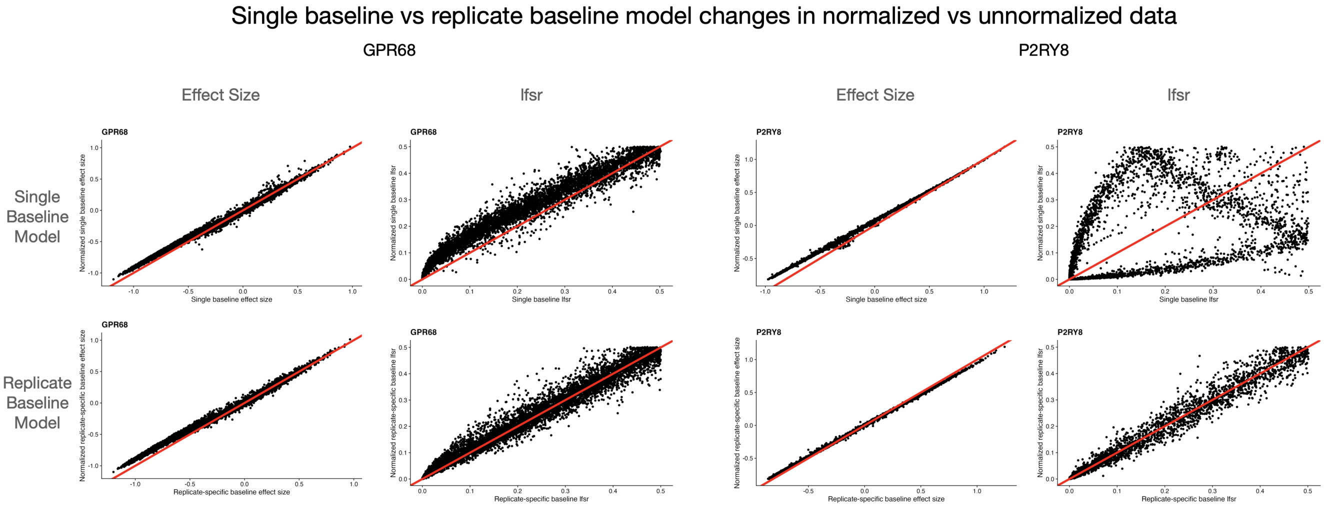


Figure 18: Side-by-side comparison of the impact of cell proportion normalization on the results of single baseline Lilace and replicate-specific baseline Lilace. The y-axis of each plot represents the model result on the normalized data and the x-axis the unnormalized data. In these datasets, using the replicate specific baseline is able to mostly recover the effect sizes and lfsrs of the normalized data without the sorting information needed for normalization. Even in the case of GPR68 where the effect sizes are still shifted by normalization, the lfsrs are corrected.

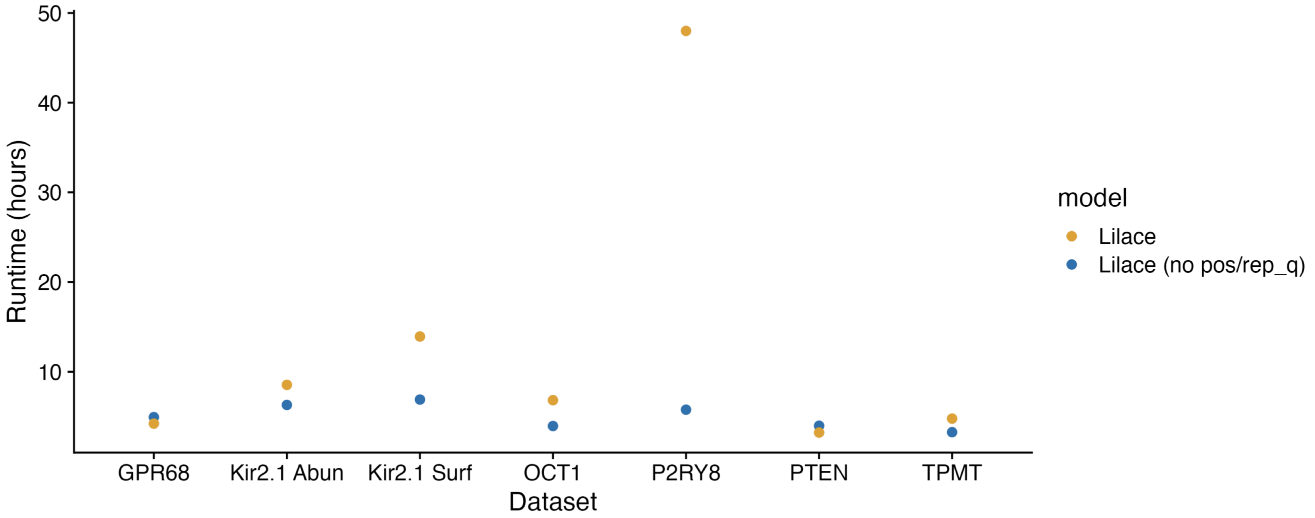


Figure 19: Runtime comparison of Lilace on different datasets on UCLA's computing cluster. As a comparison, we also plot Lilace (no pos/rep_q), which is a version of Lilace without a position effect or replicate-specific baseline parameter. For most datasets, the additional position and replicate-specific parameters do not substantially increase runtime.
